# Unraveling diel regulation of cuticle biosynthesis

**DOI:** 10.1101/2025.10.28.684701

**Authors:** Quang Ha Dang, Hyeon Jun Kim, Jeongho Choi, Da-Min Choi, Seong-Hyeon Kim, Jeong-Il Kim, Mi Chung Suh

**Affiliations:** Department of Life Science, Sogang University, Seoul 04107, Republic of Korea; Department of Integrative Food, Bioscience and Biotechnology (BK21 FOUR), Chonnam National University, Gwangju, 61186, Republic of Korea

**Keywords:** Cuticle, Cuticular wax, Cutin, Diel regulation, E3 ligase, Phytochrome, Transcription factor

## Abstract

The plant cuticle is vital for growth and adaptation to environmental stresses. Although cuticle biosynthesis is dynamically regulated by environmental cues, the molecular mechanisms integrating these external signals with intracellular responses remain largely unknown. Here, we discovered that cuticle biosynthesis is precisely orchestrated by two distinct regulatory modules acting in synchrony with the diel cycle. Daylight is perceived by phytochrome B (phyB), which suppresses cuticular wax biosynthesis by degradation of PIF4, a phytochrome-interacting bHLH factor that activates wax biosynthetic genes. This suppression is alleviated when phyB itself is degraded by the E3 ubiquitin ligase LRB, leading to diurnal activation of PIF4. In contrast, loss-of-function and transcriptional assays of *CFLAP1* demonstrated its direct negative role in cutin biosynthesis. At night, the E3 ubiquitin ligase COP1 mediates proteasomal degradation of CFLAP1, thereby promoting cutin accumulation via activation of *BDG1*. Together, these results reveal that two regulatory modules, LRB-phyB-PIF4 and COP1-CFLAP1 coordinate the diel regulation of cuticle formation, ensuring timely assembly of the protective barrier.

## INTRODUCTION

Daily light-dark cycles have driven sessile plants to adapt to diurnal environmental changes that regulate photosynthesis, flowering, and defence responses against biotic and abiotic stresses (Dodd et al., 2005; Andres and Coupland, 2012; Roeber et al.,2021). The plant cuticle, the outermost physical barrier between plants and the environment, facilitates growth, development, and tolerance to terrestrial stresses (Yeats and Rose, 2013; Ingram and Nawrath, 2017). This hydrophobic layer is composed of cutin and cuticular waxes, deposited in the epidermis of aerial tissues, root caps, and wounded surfaces (Suh et al., 2005; Berhin et al., 2019; Lee et al., 2025). Cutin serves as a fundamental structural component, forming a polyester matrix, whereas cuticular waxes are either embedded within the cutin or deposited on the surface as epicuticular crystals (Kunst and Samuels, 2003; Yeats and Rose, 2013). Polyester cutin is a polymer composed of C16- and C18-fatty acids (FAs) and their derivatives, including ω-hydroxy fatty acids (HFAs) and 1, ω-dicarboxylic acids (DCAs), which are generated by cytochrome P450 family enzymes and oxidoreductases (Li-Beisson et al., 2013). Cuticular waxes mainly consist of very long-chain fatty acids (VLCFAs) and their derivatives, including alkanes (AKs), primary and secondary alcohols (PAs and SAs), aldehydes (ALs), ketones (KEs), and wax esters (WEs), which are synthesized through alkane- and alcohol-forming pathways (Samuels et al., 2008; Bernard and Joubès, 2013; Lewandowska et al., 2020; Lee and Suh, 2022).

Light is one of the most influential environmental cues that drive diel changes in plant growth and development, including cuticle formation (Shepherd and Griffiths, 2006). Light enhances wax deposition in various plant species such as kale, swede, and Arabidopsis (Shepherd et al., 1995; Go et al., 2014). Furthermore, the biosynthesis of cuticular waxes is finely modulated by qualitative properties of light, such as light intensity and wavelength (Huang et al., 2020). Interestingly, the expression of cutin biosynthetic genes such as *BDG1* and *LACS2* is upregulated during etiolation but markedly reduced during photomorphogenesis (Ma et al., 2024). Together, these findings suggest a tight coupling between diel light signaling and cuticle metabolism. On the other hand, loss-of-function mutants defective in cutin biosynthesis, such as *lacs2, bdg1*, and *cyp86a8*, exhibit growth retardation and developmental abnormalities, including organ fusion, suggesting that cutin production is a fundamental process regulated throughout plant growth and development (Wellesen et al., 2001; Xiao et al., 2004; Tang et al., 2007). Notably, cutin—but not wax— biosynthesis is particularly enhanced in the rapidly elongating upper stem regions of Arabidopsis (Suh et al., 2005). Considering that plant elongation growth is predominantly promoted during the night (Apelt et al., 2017; Urrea-Castellanos et al., 2022), these observations raise the possibility that cutin biosynthesis may be positively associated with nocturnal growth. However, the molecular mechanisms underlying the nighttime regulation of cutin biosynthesis remain largely unexplored.

Over the past decades, extensive studies have identified a network of transcriptional regulators—including HDG1, CFLAP1, MYB96, WIN1, DEWAX, and SPL9—that control cuticle biosynthesis during development and in response to environmental stimuli such as drought and diurnal light cycles (Kannangara et al., 2007; Seo et al., 2011; Wu et al., 2011; Go et al., 2014; Li et al., 2016; Li et al., 2019). Despite these advances, the upstream signaling mechanisms that modulate the activity and stability of these transcription factors remain poorly understood. The factors directly mediating environmental cues and plant responses include photoreceptors, such as red/far-red light-sensing phytochrome B (phyB). phyB and its related E3 ubiquitin ligases, COP1, SPA1, LRB, HOS1, and BOP2 mediate light responses under continuous red light, including germination, de-etiolation, leaf expansion, inhibition of stem and petiole elongation, and flowering (Rockwell et al., 2006; Christians et al., 2012; Lu et al., 2015; Kim et al., 2017a; Zhang et al., 2017). However, how plants integrate light perception and signal transduction to coordinate the diel regulation of cutin and cuticular wax biosynthesis remains elusive. In this study, we demonstrate that plants regulate cuticle biosynthesis according to light-dark cycles via two distinct modules, LRB-phyB-PIF4 and COP1-CFLAP1, each stimulating diurnal wax and nocturnal cutin accumulation, respectively. These findings expand our insights into diel regulation of cuticle biosynthesis in Arabidopsis.

## RESULTS

### phyB negatively regulates cuticular wax biosynthesis

To investigate how plants perceive and transduce light signals to regulate cuticular wax biosynthesis, we focused on a previous report showing that maize phytochrome B (phyB) mutants exhibit increased total wax loads in leaves (Qiao et al., 2020). We therefore examined the role of phyB in Arabidopsis cuticular wax biosynthesis. The total wax load increased by approximately 20%, 60%, and 50% in the leaves of *phyA-211*, *phyB-9*, and *phyA-211 phyB-9* mutants, respectively, compared with the wild type (WT) (Figure 1A). Levels of alkanes (AKs) and primary alcohols (PAs) were significantly elevated in *phyB-9* and *phyA-211 phyB-9* (Figure 1A and Supplemental Figure 1A). Consistent with these findings in the Col-0 background, total wax amounts were higher by approximately 47% in *phyA-201 phyB-5* and lower to approximately 66% of WT levels in the *PHYB*-overexpressing (OX) line in the L*er* background (Figure 1B and Supplemental Figure 1B). Expression of wax biosynthetic genes was markedly upregulated in leaves of *phyB-9, phyA-211 phyB-9* (Col-0), and *phyA-201 phyB-5* (Ler), but downregulated in *PHYB* OX leaves (Figures 1C and 1D). In particular, transcript levels of *CER1, CER4, KCS2,* and *SOH1* were increased in *phyB* and *phyA phyB*, whereas decreased in *PHYB* OX leaves. Collectively, these results demonstrate that phyB acts as a negative regulator of cuticular wax biosynthesis in Arabidopsis, modulating wax production in response to light conditions.

**Figure 1.**
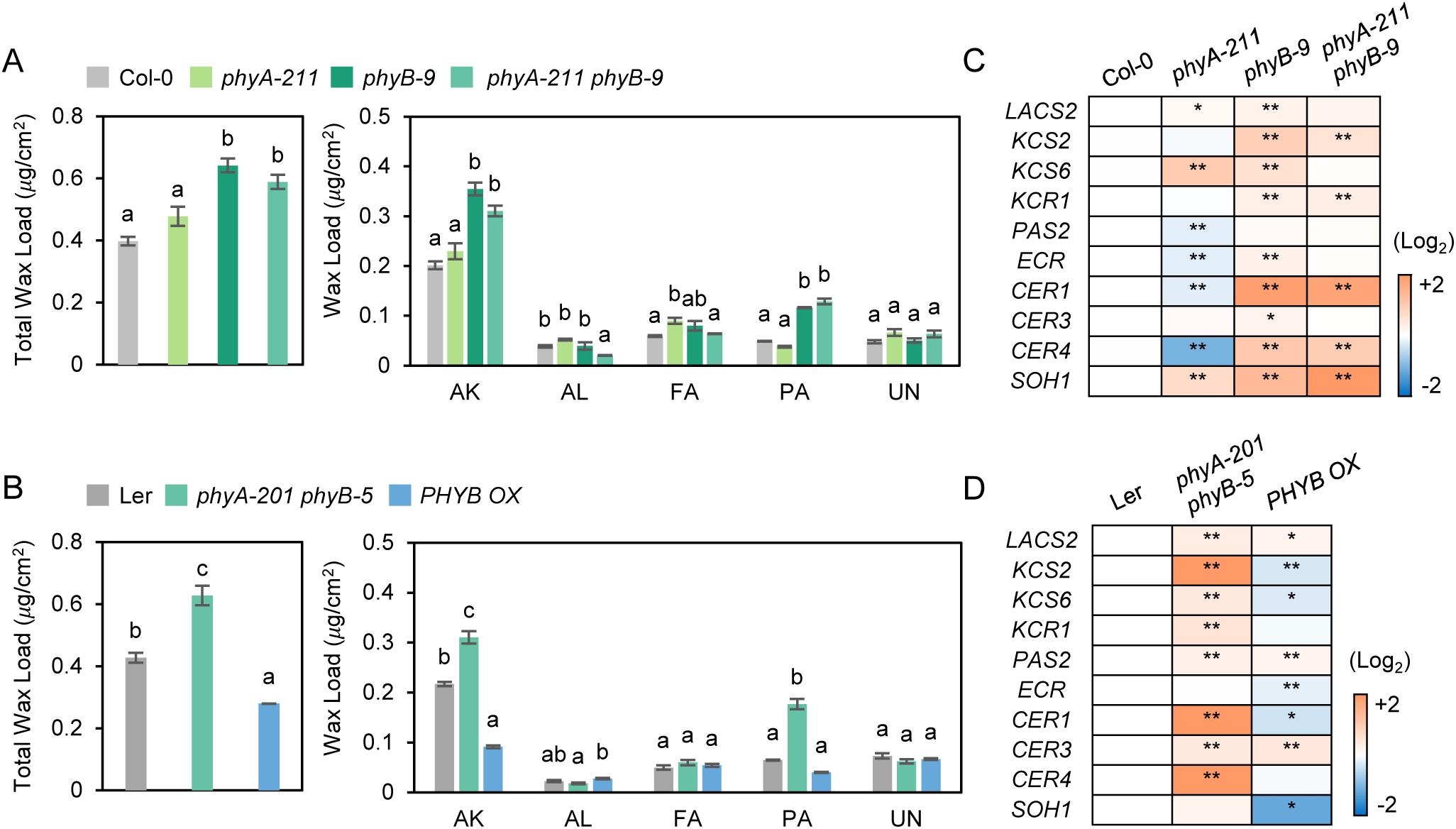
Phytochrome B negatively regulates cuticular wax biosynthesis. (A) Quantification of cuticular wax loads in 3-week-old leaves of wild type (Col-0), *phyA-211*, *phyB-9*, and *phyA-211 phyB-9*. (B) Quantification of cuticular wax loads in 3-week-old leaves of wild type (Ler), *phyA-201 phyB-5*, and *PHYB OX*. (C) Heatmap visualizing expression of cuticular wax biosynthesis-related genes in 3-week-old leaves of Col-0, *phyA-211*, *phyB-9*, and *phyA-211 phyB-9* harvested at 10:30. (D) Heatmap visualizing expression of cuticular wax biosynthesis-related genes in leaves of Ler, *phyA-201 phyB-5*, and *PHYB OX* harvested at 10:30. (A and B) Each value represents the mean ±SD of three individual replicates. AK, alkanes; AL, aldehydes; FA, fatty acids; PA, primary alcohols; UN, unidentified. Different letters indicate statistically significant differences using one-way ANOVA with Tukey’s test (*P* < 0.01). (C and D) Asterisks indicate statistically significant differences determined by Student’s t-test (*, *P* < 0.05; **, *P* < 0.01).

### PIF4 activates cuticular wax biosynthesis by directly binding to the promoters of **KCS2**, **CER1**, and **CER4**, thereby upregulating their expression

Alterations in the transcript levels of wax biosynthetic genes in *phyB* and *PHYB* OX suggest that transcription factors downstream of phyB may activate cuticular wax biosynthesis. The PIF family of transcription factors, which physically interact with phyB under light conditions (Lorrain et al., 2008), were considered as potential candidates. To identify the most relevant PIFs among the eight members in Arabidopsis, we considered two criteria: transcript abundance and protein stability during the light period. The expression levels of PIF4 and PIF5 were significantly higher than those of other PIFs during the light phase, although PIF5 expression was also detected in darkness (Supplemental Figure 2A). Consistently, PIF4 and PIF5 proteins accumulated during daytime until 14:30, but PIF5 remained detectable at night (Supplemental Figure 2B). These temporal expression patterns suggest that PIF4 is the primary mediator of light-responsive cuticular wax biosynthesis.

To test the hypothesis that PIF4 positively regulates wax biosynthesis, we analyzed cuticular wax content in the leaves of WT (Col-0), *pif4*, *pifQ* (*pif1pif3pif4pif5*), a *PIF4* complementation line in *pifQ* background (*PIF4/pifQ*), and a *PIF4* overexpression line (*PIF4*-OX). Total wax amounts were reduced by approximately 10% and 20% in *pif4* and *pifQ*, respectively, whereas they were increased by approximately 30% and 85% in *PIF4/pifQ* and *PIF4*-OX relative to the WT, respectively (Figure 2A and 2B). In particular, levels of AKs and PAs significantly decreased in *pif4* and *pifQ*, but markedly elevated in *PIF4/pifQ* and *PIF4-OX* (Figure 2A, 2B and Supplemental Figure 3). The transcript levels of *KCS2, CER1*, and *CER4*, which are involved in AK and PA biosynthesis, were significantly reduced in *pif4* and *pifQ* leaves, but increased in *PIF4/pifQ* and *PIF4-OX* leaves compared with WT (Figure 2C). No significant differences in total cutin monomer levels were detected between WT and *pif4, pifQ* or *PIF4/pifQ,* except for moderate increases in DCAs and ω-HFAs in *PIF4/pifQ* (Supplemental Figure 4). Collectively, these results demonstrate that PIF4 acts as a positive regulator of cuticular wax biosynthesis.

**Figure 2.**
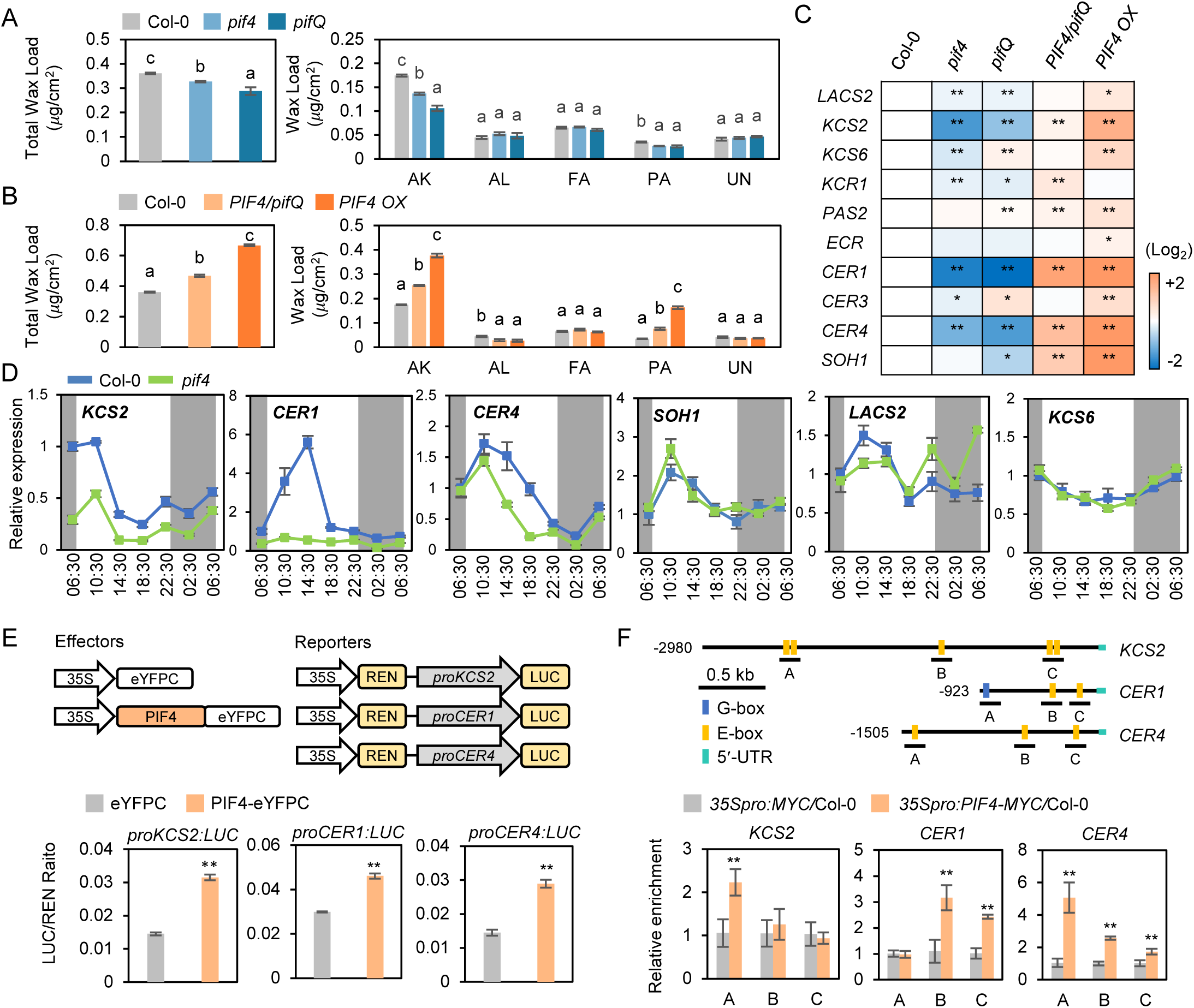
PIF4 activates cuticular wax biosynthesis by its direct binding to the promoter regions of *KCS2, CER1,* and *CER4* during the daytime. (A) Quantification of cuticular wax loads in 3-week-old leaves of wild type (Col-0), *pif4*, and *pifQ*. (B) Quantification of cuticular wax loads in 3-week-old leaves of Col-0, *PIF4pro:PIF4-MYC/pifQ* (*PIF4/pifQ*), and *35Spro:PIF4-MYC*/Col-0 (*PIF4 OX*). (C) Heatmap showing expression of cuticular wax biosynthesis-related genes in 3-week-old leaves of Col-0, *pif4*, *pifQ*, *PIF4/pifQ*, and *PIF4 OX* harvested at 10:30. (D) Diurnal expression patterns of *KCS2*, *CER1*, *CER4*, *SOH1*, *LACS2* and *KCS6* in Col-0 and *pif4.* Leaves of 3-week-old plants grown under long-day conditions were harvested at indicated time points. Transcript levels were examined by RT–qPCR. (E) Dual-luciferase assays were performed in *N. benthamiana* to examine transcriptional activities of PIF4-eYFPC on the promoter regions of *KCS2, CER1,* and *CER4.* LUC activity values were normalized to *Renilla* (REN) luciferase activity to represent relative promoter activities. (F) ChIP-qPCR assays showed that PIF4 associates with cis-elements within the promoters of *KCS2*, *CER1* and *CER4 in vivo*. UTR, Untranslated region. (A, B, D, E, and F) Values represent mean ±SD from three replicate experiments. Different letters indicate statistically significant differences using one-way ANOVA with Tukey’s test (*P* < 0.01). AK, alkanes; AL, aldehydes; FA, fatty acids; PA, primary alcohols; UN, unidentified. (C, E, and F) Asterisks indicate statistically significant differences determined by Student’s t-test (*, *P* < 0.05; **, *P* < 0.01).

To identify target genes of PIF4 involved in AK and PA biosynthesis, we performed RT-qPCR analysis in WT and *pif4* leaves over the diel cycle. In WT, *KCS2, CER1, CER4, SOH1*, and *LACS2* displayed typical diurnal expression patterns, peaking at 10:30 or 14:30 during the light period and declining thereafter. In contrast, *KCS6* exhibited only minor variation during the day, showing a modest decrease during the light phase followed by a slight recovery (Figure 2D). In *pif4* mutant, transcript levels of *KCS2, CER1,* and *CER4* were noticeably reduced compared with WT, whereas no significant differences were observed for *SOH1*, *LACS2,* and *KCS6* (Figure 2D). Notably, the diurnal oscillation of *CER1* expression was completely diminished in *pif4* leaves. Therefore, these findings indicate that PIF4 regulates the transcription of cuticular wax biosynthetic genes in a diurnal manner. We next examined whether PIF4 directly activates the expression of *KCS2, CER1,* and *CER4*. Transient dual-luciferase reporter assays in tobacco (*Nicotiana benthamiana*) leaves revealed that LUC reporter activities driven by the promoters of *KCS2, CER1,* and *CER4* were significantly enhanced in the presence of PIF4 compared with the control (Figure 2E). Consistently, chromatin immunoprecipitation (ChIP) assays demonstrated that PIF4 directly binds to the promoter regions of *KCS2* (region A), *CER1* (regions B and C), and *CER4* (regions A, B, and C), each containing a canonical E-box motif (Figure 2F).

### Disruption of **phyB** stabilizes PIF4, leading to upregulation of cuticular wax biosynthesis during the daytime

We next examined whether phyB functions upstream of PIF4 to inhibit upregulation of cuticular wax biosynthesis. Immunoblot analysis revealed that PIF4 protein levels were markedly higher in *phyB-9* than in WT (Figure 3A). Consistent with the elevated PIF4 levels, the expression of its direct target genes, *KCS2, CER1*, and *CER4*, was significantly upregulated in *phyB-9* compared with WT at 10:30 and/or 14:30, with *SOH1* and *KCS6* transcripts likewise increased (Figure 3B). Given that light-activated phyB represses PIF4 activity (Lorrain et al., 2008), wax biosynthesis is suppressed while phyB remains active. The enhanced wax production observed under light therefore indicates that phyB must be degraded to relieve PIF4 from repression.

**Figure 3.**
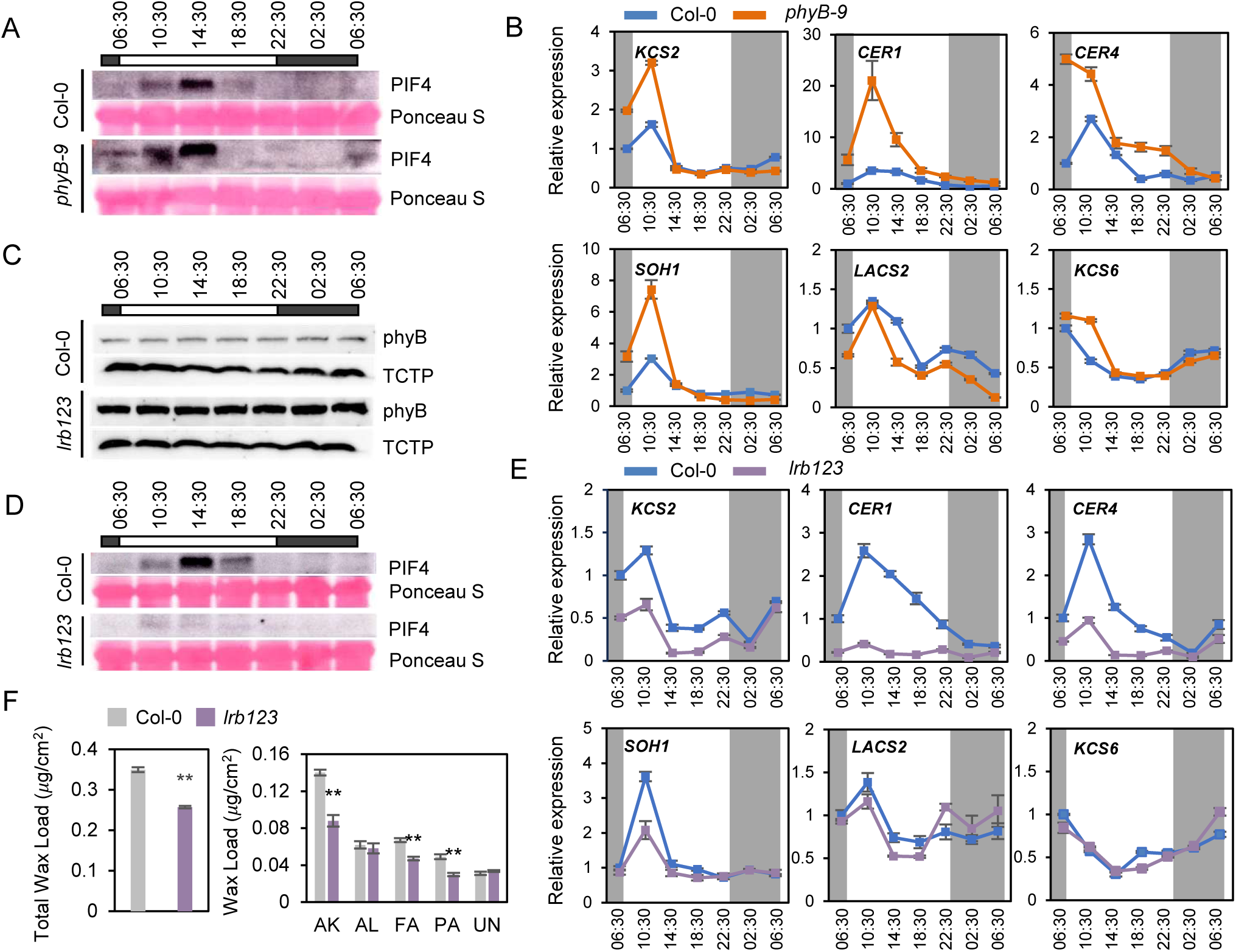
LRB-mediated destabilization of phyB results in PIF4 stabilization, thereby promoting cuticular wax biosynthesis during the daytime. (A) Immunoblot analysis of PIF4 protein in 3-week-old wild type (Col-0) and *phyB-9* using an anti-PIF4 antibody. (B) Diurnal expression patterns of *KCS2*, *CER1*, *CER4*, *SOH1*, *LACS2* and *KCS6* in Col-0 and *phyB-9.* (C) Immunoblot analysis of phyB protein in 2-week-old Col-0 and *lrb123* plants using an anti-phyB antibody. Total protein loading was assessed by anti-TCTP antibody. 2-week-old plants were harvested at indicated time points. (D) Immunoblot analysis of PIF4 protein in 3-week-old Col-0 and *lrb123* using an anti-PIF4 antibody. (E) Diurnal expression patterns of *KCS2*, *CER1*, *CER4*, *SOH1*, *LACS2* and *KCS6* in Col-0 and *lrb123.* (F) Quantification of cuticular wax loads in 3-week-old leaves of Col-0 and *lrb123*. Values represent mean ±SD of three replicate experiments. Asterisks indicate statistically significant differences determined by Student’s t-test (*, *P* < 0.05; **, *P* < 0.01). AK, alkanes; AL, aldehydes; FA, fatty acids; PA, primary alcohols; UN, unidentified. (A, B, D, and E) Leaves of 3-week-old plants were harvested at indicated time points. (A and D) Total protein loading was assessed by Ponceau S staining. (B and E) Transcript levels were examined by RT–qPCR. Values represent mean ±SD of three replicate experiments.

Because Light-Responsive BTB (LRB) proteins, adaptors of E3 ubiquitin ligase complexes, promote the degradation of active phyB (Pfr form) together with PIF3 (Ni et al., 2014), we examined phyB and PIF4 abundance in WT and the *lrb1lrb2-2lrb3* (*lrb123*) triple mutant leaves. In *lrb123*, phyB protein levels were substantially increased, whereas PIF4 protein levels were reduced compared with WT (Figure 3C and 3D). In line with the lowered levels of PIF4, daytime expression of *KCS2, CER1*, *CER4,* and *SOH1* were significantly downregulated in *lrb123* relative to WT, while the expression of *LACS2* and *KCS6* remained unchanged (Figure 3E). Cuticular wax analysis further showed that total wax loads in *lrb123* were reduced to approximately 60% of WT levels (Figure 3F). Significant decreases in the levels of AKs, VLCFAs, and PAs, the major wax components, were detected in *lrb123* leaves relative to WT (Figure 3F and Supplemental Figure 5). These results indicate that LRB-dependent phyB degradation is important for the upregulation of cuticular wax biosynthesis under light conditions.

### CFLAP1 negatively regulates cutin biosynthesis

Given that the cuticular structure where cutin is deposited underneath the wax layer (Yeats and Rose, 2013), cuticular wax accumulation during the day should be preceded by cutin deposition. Suh et al. (2005) reported that in Arabidopsis stems, cutin monomer loads were approximately 2-fold higher in the elongating upper region than in the basal region, whereas total wax loads showed no significant difference. Comparative analysis of cuticle biosynthetic gene expression between the upper and lower stem regions revealed that cutin-related genes were generally more strongly upregulated than wax-related genes (Suh et al., 2005; Supplemental Figure 6A), suggesting the crucial role of cutin biosynthesis in highly expanding tissues. Considering that plant elongation growth primarily occurs at night (Apelt et al., 2017), we hypothesized that cutin biosynthesis is closely related with nocturnal growth. Supporting this idea, microarray data from Blaesing et al. (2005) showed that most cutin biosynthetic genes were upregulated under dark conditions (Supplemental Figure 6B). Among the transcription factors implicated in cuticle formation (Lee and Suh, 2022), the expression of the negative regulator *CFLAP1* (Li et al., 2016) was markedly decreased in Arabidopsis leaves in darkness relative to light, suggesting that darkness-induced suppression of *CFLAP1* may contribute to the enhanced expression of cutin biosynthetic genes.

To examine the function of CFLAP1 in cuticle development, we isolated two loss-of-function mutants, *cflap1-1* and *cflap1-2* (Supplemental Figure 6C and 6D), and analyzed cutin and wax composition and amounts by gas chromatography. Total cutin monomer content, particularly that of ω-HFAs and DCAs, was significantly increased in *cflap1* mutant leaves compared with WT, whereas cuticular wax content showed no significant change (Figure 4A, 4B and Supplemental Figure 7A and 7B). These results indicate that CFLAP1 acts as a negative regulator in cutin deposition.

**Figure 4.**
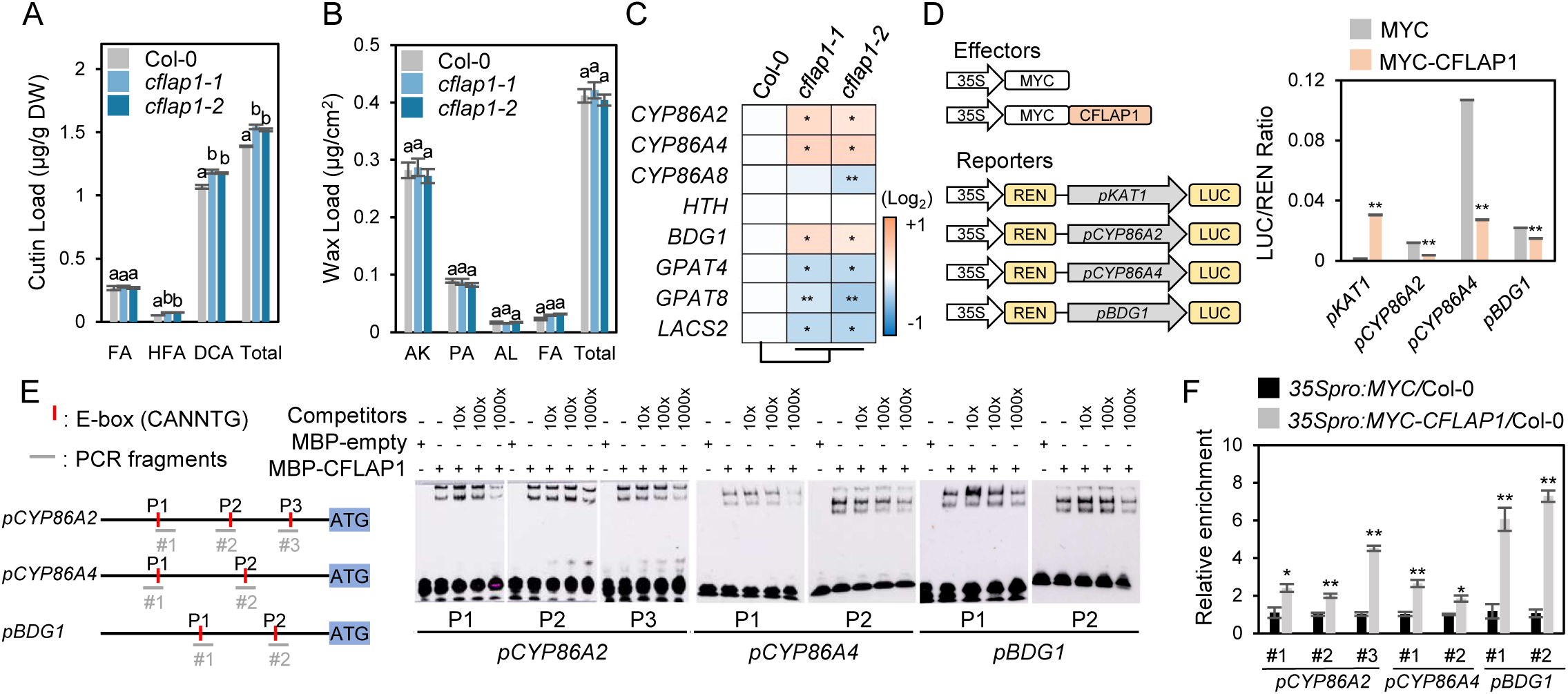
CFLAP1 negatively regulates cutin biosynthesis by its direct binding to the promoter regions of *CYP86A2*, *CYP86A4* and *BDG1.* (A and B) Quantification of cutin monomer (A) and cuticular wax (B) loads in 3-week-old leaves of wild type (Col-0), *cflap1-1*, and *cflap1-2*. Each value represents the mean ±SD of three individual replicates. FA, fatty acids; HFA, ⍵*-*hydroxy fatty acids; DCA, dicarboxylic acids; AK, alkanes; PA, primary alcohols; AL, aldehydes. Different letters indicate statistically significant differences using one-way ANOVA with Tukey’s test (*P* < 0.01). (C) Relative expression levels of cutin biosynthetic genes in 12-day-old seedlings of Col-0, *cflap1-1*, and *cflap1-2* harvested at 18:30 are visualized as a heatmap. Scale bar, log_2_ fold change. (D) Dual-luciferase assays were performed in *N. benthamiana* to examine transcriptional activities of MYC-CFLAP1 on the promoter regions of *KAT1, CYP86A2, CYP86A4,* and *BDG1.* LUC activity values were normalized to *Renilla* (REN) luciferase activity to represent relative promoter activities. Data represent means ±SD from three biological replicates. (E) EMSA assay showing binding of recombinant CFLAP1 protein to the promoters of *CYP86A2* (P1, P2, and P3), *CYP86A4* (P1 and P2), and *BDG1* (P1 and P2), and competition of binding with increasing concentration of cold DNA probes. Diagram depicts the promoters of *pCYP86A2, pCYP86A4,* and *pBDG1* with putative binding E-box motifs. (F) ChIP-qPCR assays showed that CFLAP1 associates with cis-elements within the promoters of *CYP86A2*, *CYP86A4* and *BDG1 in vivo*. Data are means ±SD (n = 3). (D and F) Asterisks indicate statistically significant differences determined by Student’s t-test (*, *P* < 0.05; **, *P* < 0.01).

Based on the marked upregulation of cutin biosynthetic genes in response to darkness, when nocturnal growth occurs, we examined whether *CFLAP1* transcripts and protein levels are diurnally regulated. *CFLAP1* mRNA levels increased during the light period, showing two distinct peaks at 10:30 and 18:30, and declined significantly at night (Supplemental Figure 8A and 8B). In Arabidopsis seedlings overexpressing *CFLAP1* fused with a MYC epitope under the control of CaMV 35S promoter (*CFLAP1* OX), the accumulation pattern of MYC-CFLAP1 closely paralleled with that of *CFLAP1* mRNA (Supplemental Figure 8C), indicating that CFLAP1 expression is subject to diurnal regulation.

To elucidate the regulatory role of CFLAP1 in cutin biosynthesis, we analyzed the expression of cutin biosynthetic genes in WT, *cflap1-1*, and *cflap1-2* leaves at 18:30, when *CFLAP1* expression peaks. The transcript levels of *CYP86A8, GPAT4, GPAT8*, and *LACS2* were reduced, whereas those of *CYP86A2, CYP86A4*, and *BDG1* were significantly increased in *cflap1* mutants relative to WT (Figure 4C), suggesting that *CYP86A2, CYP86A4,* and *BDG1* are potential downstream targets of CFLAP1. Because CFLAP1 is a bHLH transcription factor that binds to E-box motif (5′-CANNTG-3′), approximately 3-kb promoter regions of *CYP86A2, CYP86A4* and *BDG1* including E-box motifs were used for the generation of reporter constructs. The effector construct contained *MYC-CFLAP1* driven by the CaMV 35S promoter. In a dual-luciferase assay, MYC-CFLAP1 significantly repressed the LUC/REN activity driven by the *pBDG1, pCYP86A2* and *pCYP86A4* promoters by approximately 1.5- to 4-fold compared with the control (Figure 4D). To test direct binding, electrophoretic mobility shift assays (EMSAs) were performed using recombinant maltose-binding protein (MBP)–CFLAP1 purified from *E. coli* (Supplemental Figure 9) and promoter fragments from *pCYP86A2, pCYP86A4*, and *pBDG1*, each containing one or more E-boxes (Figure 4E). EMSAs demonstrated that MBP-CFLAP1 specifically interacts with these promoter fragments (Figure 4E). Consistently, ChIP-qPCR assay showed enrichment of CFLAP1 on the E-box regions of *CYP86A2*, *CYP86A4,* and *BDG1* promoters in *MYC-CFLAP1* OX seedlings relative to the control (Figure 4F). These results indicate that CFLAP1 directly binds to the E-box in the promoters of *CYP86A2, CYP86A4*, and *BDG1*, thereby repressing their transcription to negatively regulate cutin biosynthesis.

### COP1 interacts with and degrades CFLAP1 proteins via 26S proteasome pathway

Because CFLAP1 protein levels were markedly lower at night than those during the day (Supplemental Figure 8C), we investigated whether COP1, which acts in the nucleus under dark conditions, is involved in regulating CFLAP1 stability. Luciferase complementation imaging (LCI) assays showed a physical interaction between COP1 and CFLAP1 when co-expressed in tobacco leaves (Figure 5A). Bimolecular fluorescence complementation (BiFC) analysis further showed nuclear YFP signals in tobacco epidermal cells coexpressing *COP1-eYFPN* and *CFLAP1-eYFPC* (Figure 5B). Co-immunoprecipitation (Co-IP) assays confirmed this interaction: COP1-HA proteins were detected in extracts from tobacco leaves co-expressing *MYC-CFLAP1* and *COP1-HA* and immunoprecipitated with anti-MYC antibody, but not in extracts co-expressing with empty *MYC* and *COP1-HA* (Figure 5C). Immunoblot analyses demonstrated that the COP1–CFLAP1 interaction promotes CFLAP1 ubiquitination and subsequent 26S proteasome-mediated degradation, as indicated by the increased accumulation of ubiquitinated CFLAP1 following MG132 treatment (Figure 5D and 5E). To examine COP1-dependent degradation *in planta*, *MYC-CFLAP1* OX lines were crossed with the *cop1-4* mutant. Immunoblot analysis revealed markedly higher levels of MYC-CFLAP1 protein in the *cop1-4* background than in WT (Col-0), whereas transcript levels of *MYC-CFLAP1* were comparable between the two genotypes (Figure 5F). These results demonstrate that COP1 mediates ubiquitination and 26S proteasome-dependent degradation of CFLAP1.

**Figure 5.**
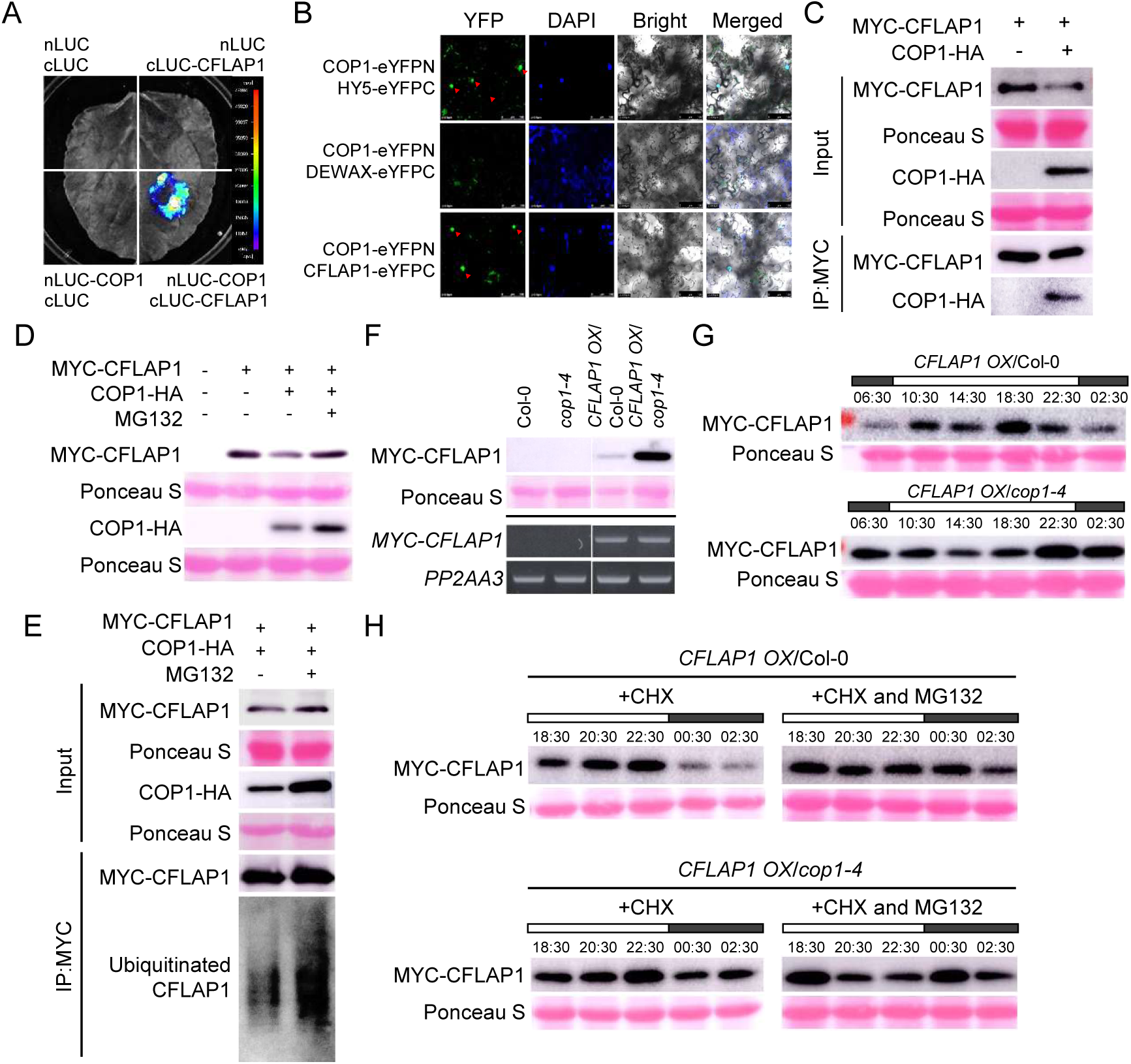
COP1 interacts with CFLAP1 and facilitates its ubiquitination and degradation via 26S proteasome system during the nighttime. (A) LCI assay of interaction between nLUC-COP1 and cLUC-CFLAP1 *in vivo*. (B) BiFC assay of nuclear interaction between COP1-eYFPN and CFLAP1-eYFPC. YFP fluorescence (green) indicates physical interaction between COP1 and each target protein. DAPI staining (blue) marks nuclei. Scale bars, 100 µm. (C) Co-immunoprecipitation (Co-IP) of the interaction between COP1-HA and MYC-CFLAP1. Proteins were immunoprecipitated with an anti-MYC (α-MYC) antibody and detected using α-MYC and α-HA antibodies. (D) Immunoblot analysis of the co-expression of MYC-CFLAP1 with COP1-HA with the proteasome inhibitor MG132. Proteins were detected using α-MYC and α-HA antibodies. (E) Ubiquitination assay of MYC-CFLAP1 upon co-expression with COP1-HA within 100 μM MG132 treatment. Proteins were immunoprecipitated with an anti-MYC (α-MYC) antibody and detected using α-MYC and anti-Ubiquitin (α-Ub) antibodies. (F) Immunoblot (top) and RT-PCR (bottom) of *MYC-CFLAP1* expression in *Arabidopsis* CFLAP1-overexpressing (*CFLAP1 OX*) lines in the wild type (Col-0) or *cop1-4* background. (G) Diurnal accumulation patterns of CFLAP1 protein in *CFLAP1 OX*/Col-0 and *CFLAP1 OX*/*cop1-4* seedlings collected at various time points. (H) 100 μM cycloheximide (CHX) and 100 μM CHX + 100 μM MG132 treatment assays of CFLAP1 protein stability in *CFLAP1 OX*/Col-0 and *CFLAP1 OX*/*cop1-4* seedlings at various time points. (F-H) Proteins were detected using α-MYC. (C-H) Ponceau S staining indicates equal protein loading.

Next, we examined whether CFLAP1 protein levels depend on COP1 activity and subcellular localization in *CaMV35S:MYC-CFLAP1/*WT and *CaMV35S:MYC-CFLAP1/cop1-4* seedlings across the day-night cycle. In the WT background, MYC-CFLAP1 levels peaked during the daytime and declined rapidly at night. By contrast, in *cop1-4* background, MYC– CFLAP1 proteins accumulated continuously throughout all examined time points, with no decrease detected at night (Figure 5G). To further examine whether COP1 mediates nocturnal degradation of CFLAP1 via the 26S proteasome pathway, seedlings were treated with cycloheximide (CHX) in the presence or absence of MG132. In *CFLAP1* OX/Col-0, degradation of MYC-CFLAP1 at night under CHX treatment was abolished by MG132, whereas MYC-CFLAP1 levels remained largely unchanged in *CFLAP1 OX*/*cop1-4* under both CHX and CHX + MG132 treatments (Figure 5H). These results indicate that COP1 mediates the nocturnal ubiquitination and subsequent 26S proteasome-dependent degradation of CFLAP1.

### The COP1-CFLAP1 module positively regulates cutin biosynthesis during nighttime

COP1-mediated degradation of CFLAP1 prompted examination of cuticle phenotypes in two loss-of-function *cop1* alleles (*cop1-4* and *cop1-6*). Cuticle permeability to toluidine blue O (TBO) was dramatically increased in leaves of *cop1* mutants compared with WT (Figure 6A). Transmission electron microscopy (TEM) analysis revealed a reduced cuticular proper layer and thinner cell wall and cuticle in *cop1-4* relative to WT (Figure 6B). Consistent with these structural defects, cuticular transpiration and chlorophyll leaching assays showed enhanced water loss and faster diffusion of chlorophylls into 80% ethanol from leaves of *cop1-4* and *cop1-6* mutants compared with WT (Figure 6C and 6D). GC-FID analysis further revealed that leaves of *cop1-4* and *cop1-6* mutants exhibited a marked decrease in total cutin monomer levels, particularly ω-HFAs and DCAs, whereas total wax composition and content were not significantly altered (Supplemental Figure 10). These results demonstrate that COP1 is essential for normal cuticle formation by maintaining proper cutin deposition.

**Figure 6.**
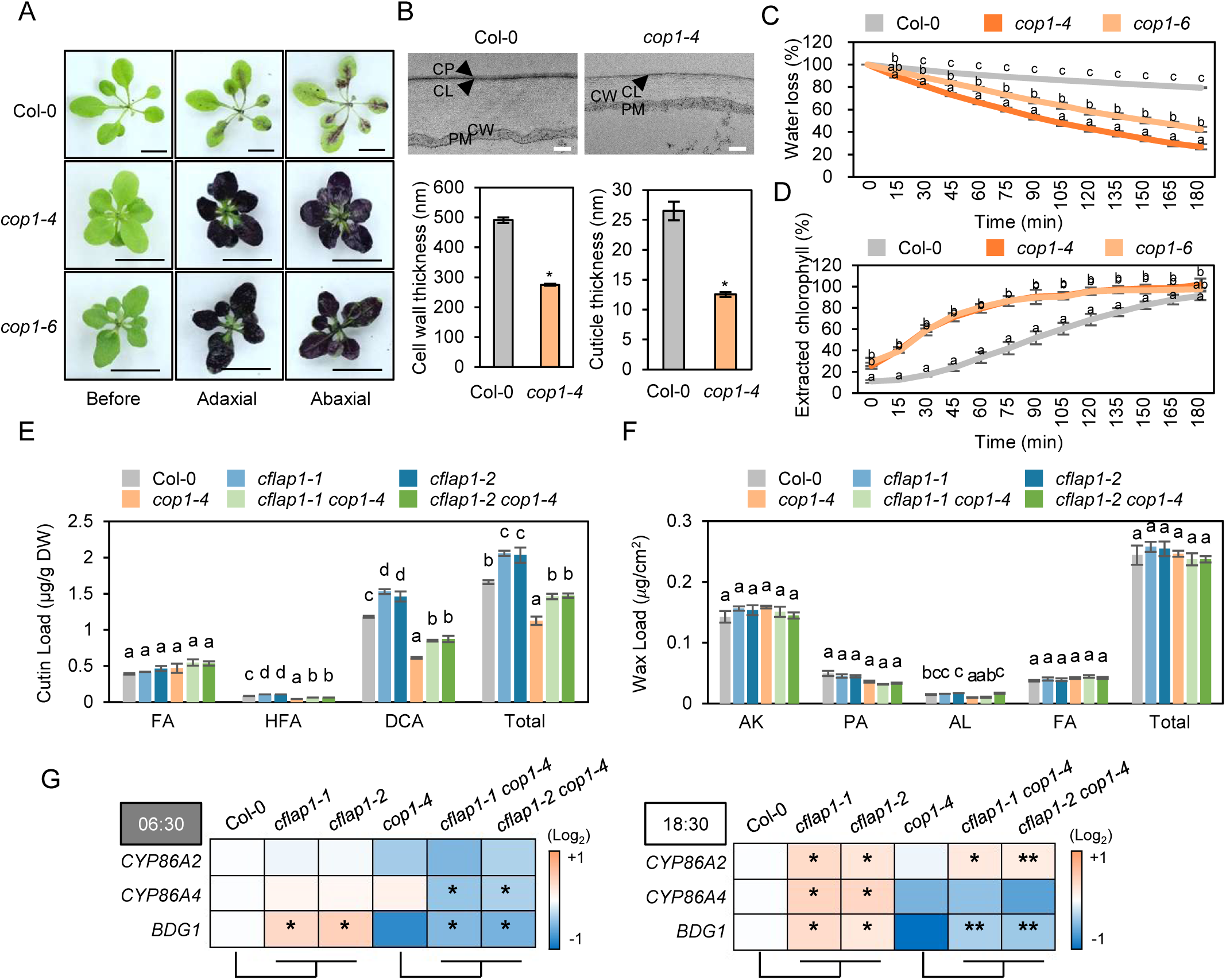
The COP1-CFLAP1 module upregulates cutin biosynthesis during the nighttime. (A) Representative images of 3-week-old rosettes from Col-0, *cop1-4* and *cop1-6* before staining, and adaxial or abaxial leaf surfaces after staining with 0.1% TBO containing 0.01% Tween 20 for 3 min. Scale bars, 1 cm. (B) Transmission electron microscopy (TEM) images of cuticle ultrastructure in the adaxial epidermis of 3-week-old leaves. The cell wall and cuticle thickness were measured using ImageJ at multiple positions. Values represent the mean ±SD of 15 (cell wall) and 20 (cuticle) measurements. CP, cuticle proper; CL, cuticular layer; CW, cell wall; PM, plasma membrane. Scale bars, 200 nm. (C) Cuticular transpiration assay showing time courses of water loss (%) in 3-week-old Col-0, *cop1-4* and *cop1-6* leaves. Values represent the mean ±SD of 3 individual replicates. (D) Chlorophyll leaching assay showing time courses of chlorophyll extraction (%) from 3-week-old Col-0, *cop1-4* and *cop1-6* leaves. Values represent the mean ±SD of 3 individual replicates. (E and F) Quantification of cutin monomer (E) and cuticular wax (F) loads in 3-week-old rosette leaves wild type (Col-0), *cflap1-1, cflap1-2, cflap1-1 cop1-4,* and *cflap1-2 cop1-4*. Values represent the mean ±SD of 3 individual replicates. FA, fatty acids; HFA, ⍵*-*hydroxy fatty acids; DCA, dicarboxylic acids; AK, alkanes; PA, primary alcohols; AL, aldehydes. (C-F) Different letters indicate statistically significant differences using one-way ANOVA with Tukey’s test (*P* < 0.01). (G) Heatmap showing relative expression levels of *CYP86A2, CYP86A4* and *BDG1* in 12-day-old seedlings of Col-0, *cflap1-1, cflap1-2, cflap1-1 cop1-4,* and *cflap1-2 cop1-4* harvested at 06:30 and 18:30. Scale bar, log_2_ fold change. (B and G) Asterisks indicate statistically significant differences determined by Student’s t-test (*, *P* < 0.05; **, *P* < 0.01).

To investigate the functional relationship between COP1 and CFLAP1 in cuticle formation, *cflap1-1* and *cflap1-2* were crossed with *cop1-4* (Supplemental Figure 11) and the resulting *cflap1-1 cop1-4* and *cflap1-2 cop1-4* double knockout mutants were analyzed for cutin monomer and cuticular wax composition and amounts by GC–FID. Leaves of both double mutants exhibited partial restoration of the reduced ω-HFA and DCA contents observed in *cop1-4* (Figure 6E and Supplemental Figure 12A). Total wax load and composition were largely unchanged, except for minor differences in C28–C30 VLCFAs and C30 PAs (Figure 6F and Supplemental Figure 12B). We next examined the transcript levels of *CYP86A2, CYP86A4* and *BDG1* identified as direct CFLAP1 target genes in WT, *cflap1-1*, *cflap1-2, cop1-4*, *cflap1-1 cop1-4,* and *cflap1-2 cop1-4* seedlings at night (06:30) and during the day (18:30). Transcript levels of *BDG1* were markedly elevated in *cflap1-1* and *cflap1-2* and the downregulation of *BDG1* observed in *cop1-4* was substantially restored in *cflap1-1 cop1-4* and *cflap1-2 cop1-4* mutants at both time points (Figure 6G). Expression of *CYP86A2* and *CYP86A4* did not show consistent patterns across the mutant lines (Figure 6G). Together, these results indicate that elevated CFLAP1 levels in *cop1* mutants primarily cause cuticle defects through repression of *BDG1*, indicating that the COP1-CFLAP1 regulatory module coordinates nocturnal activation of cutin biosynthesis.

## DISCUSSION

The plant cuticle is a crucial surface barrier that supports growth and confers tolerance to external stresses. Cuticle-defective mutants display pronounced developmental retardation and reduced stress resistance, underscoring its protective function (Wellesen et al., 2001; Xiao et al., 2004; Tang et al., 2007; Seo et al., 2011). To maintain appropriate surface properties, plants must temporally regulate cuticle biosynthesis. Our study reveals an intricate diel regulation of cuticle biosynthesis mediated by two distinct modules, the LRB-phyB-PIF4 and COP1-CFLAP1 pathways. Light-perceiving phyB inhibits PIF4 activity, which promotes cuticular wax biosynthesis by direct upregulation *of KCS2, CER1*, and *CER4*. LRB destabilizes phyB during the daytime, leading to wax biosynthesis through active PIF4. CFLAP1 negatively controls cutin biosynthesis by repressing *BDG1* expression. COP1 is nuclear-localized at night and gains access to CFLAP1, inducing its degradation via the 26S proteasomal pathway and consequently upregulating cutin biosynthesis. Nocturnal stimulation of cutin formation is followed by diurnal enhancement of cuticular wax biosynthesis. Structurally, the cuticle consists of a polyester cutin matrix associated with the cell wall, which is subsequently complemented by an outer wax layer (Yeats and Rose, 2013). Given that cell expansion often peaks at night (Urrea-Castellanos et al., 2022), we propose that rapid nocturnal cutin deposition safeguards newly exposed cell surfaces, while daytime wax biosynthesis further reinforces the barrier (Figure 7). Our findings provide a mechanistic framework for cuticle development through time-of-day coordination.

**Figure 7.**
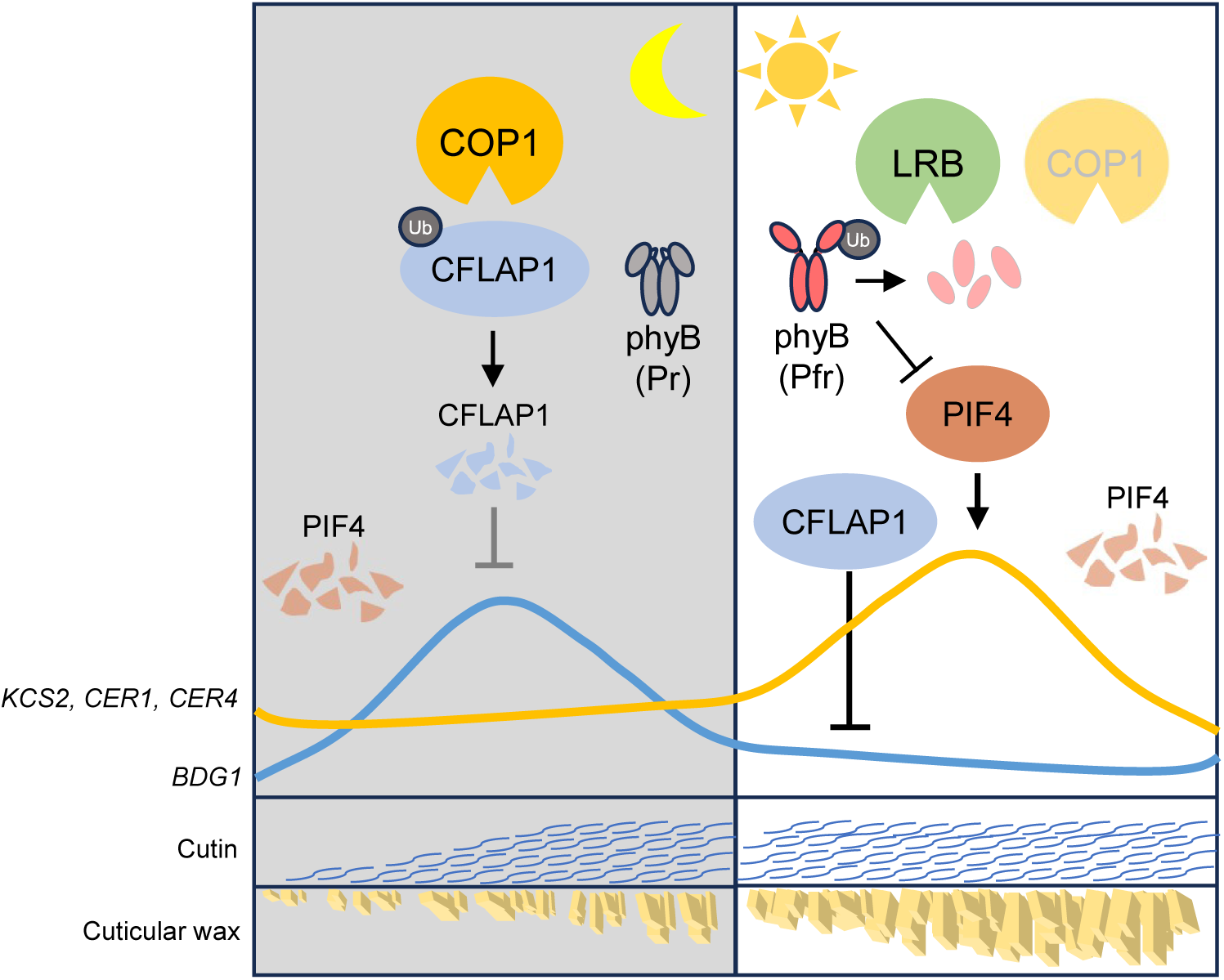
Diel regulation of cutin and cuticular wax biosynthesis. At night, nuclear-localized COP1 ubiquitinates CFLAP1, facilitating its degradation via the 26S proteasome system. Repression of CFLAP1 leads to enhanced cutin biosynthesis. During the day, light activates phyB to biologically active Pfr form. Photoactivated phyB is ubiquitinated and degraded by LRB proteins. LRB-driven depletion of active phyB releases phyB-mediated degradation of PIF4, allowing PIF4 accumulation. Accumulated PIF4 directly activates cuticular wax biosynthesis.

### LRB-phyB-PIF4 module upregulates cuticular wax biosynthesis during the daytime

Diurnal regulation of wax biosynthesis has been widely investigated, particularly at transcriptional levels. Two physically interacting transcription factors, SPL9 and DEWAX, act antagonistically to regulate *CER1* expression, optimizing wax biosynthesis during daily light-dark cycles (Go et al., 2014; Li et al., 2019). Here, we further identified PIF4 as a transcriptional activator of cuticular wax biosynthesis. PIF4 directly promotes the expression of cuticular wax biosynthetic genes *KCS2, CER1*, and *CER4* during the daytime (Figure 2D-2F), highlighting *KCS2* and *CER4* as additional diurnally regulated wax-related genes. CER1 encodes an aldehyde decarbonylase that produces AKs whereas CER4 encodes a fatty acyl-CoA reductase that converts acyl-CoA into PAs. AKs function as hydrophobic barriers that enhance drought tolerance by reducing cuticular permeability, whereas PAs increase cuticle permeability to facilitate evaporative cooling under heat stress, revealing a dynamic balance between the two pathways for environmental adaptation (Li et al., 2025). PIF4 thus acts as a critical regulator fine-tuning wax biosynthesis by contributing to the expression of both genes. Although *SOH1* expression does not appear to be regulated by PIF4 (Figure 2D), *pifQ* mutant displayed significantly reduced *SOH1* transcript levels (Figure 2C), accompanied by lower cuticular wax amounts than the *pif4* single mutant (Figure 2A). PIFs are known to act both redundantly and independently as key integrators of plant light signal responses (Leivar and Monte, 2014). Thus, in addition to PIF4, other PIFs may also contribute to the regulation of cuticular wax biosynthesis. PIF5 accumulates both transcriptionally and translationally during the daytime, in parallel with PIF4 (Supplemental Figure 2), which further supports its potential additive role in PIF4-mediated regulation of cuticular wax biosynthesis.

Despite advances in transcriptional regulation of diurnal wax accumulation, how upstream light signals are transduced to control the expression of wax biosynthetic genes remains poorly understood. Our findings demonstrate that defective phyB results in increased PIF4 protein abundance and enhanced expression of its target genes (Figure 3A and 3B). These results identify phyB as a principal light transducer that negatively regulates wax activator PIF4, likely by promoting its phosphorylation-dependent proteasomal degradation (Huq and Quail, 2002; Park et al., 2018). Among the five phytochromes (phyA-E) in Arabidopsis, phyA is a light-unstable phytochrome, unlike the other light-stable phytochromes (Pratt, 1995). Minor differences in cuticular wax amounts between *phyA-201* and wild type (Figure 1A) may reflect insufficient phyA protein abundance in wild-type plants. The *lrb123* mutant exhibits marked accumulation of phyB accompanied by reduced expression of wax biosynthetic genes and decreased wax deposition (Figure 3E and 3F). Since precise quantitative regulation of phyB abundance is required to maintain proper wax production, LRB acts as a direct repressor of phyB to ensure its optimal level. Although COP1 was identified as another E3 ligase targeting phyB (Jang et al., 2010), phyB abundance changes little during the nighttime in WT (Figure 3C). A likely explanation is that during light period, phyB is imported into the nucleus whereas COP1 is exported to the cytosol, limiting COP1 access to phyB (von Arnim and Deng, 1994; Yamaguchi et al., 1999). Accordingly, wax biosynthesis is only modestly affected in *cop1* mutants, presumably because phyB levels remain unchanged (Figure 6F and Supplemental figure 10B). Although LRB is indispensable for phyB degradation, PIF4 turnover appears to involve LRB-independent mechanisms, as its protein levels were not significantly affected in *lrb123* (Figure 3D). Under red light, BLADE-ON-PETIOLE1 (BOP1) and BOP2 assemble CUL3^BOP1/BOP2^ E3 ligases that target PIF4 for ubiquitination (Zhang et al., 2017). It would be intriguing to investigate whether BOPs affect cuticular wax biosynthesis through the regulation of PIF4 stability. Previous studies have shown that most terrestrial plants possess multiple functional LRB proteins that cooperatively regulate phytochrome-mediated light signaling (Christians et al., 2012). This evolutionary conservation suggests that LRB-dependent regulation of phyB abundance may also represent a fundamental mechanism controlling cuticular wax biosynthesis in land plants.

### The COP1-CFLAP1 module activates cutin biosynthesis at night

Previous studies have identified CFLAP1, also known as ABA-RESPONSIVE KINASE SUBSTRATE1 (AKS1), as a regulator of stomatal opening (Takahashi et al., 2017). Under high ABA conditions, SnRK2.6/OST1 phosphorylates AKS1, preventing its binding to the *KAT1* promoter and thereby suppressing potassium channel accumulation in guard cells, which leads to stomatal closure (Takahashi et al., 2017). Our research further uncovered a distinct role of CFLAP1 in regulating cutin biosynthesis. CFLAP1 acts as a transcriptional repressor of cutin related genes by directly binding to their promoters (Figure 4). At night, COP1-mediated degradation of CFLAP1 (Figure 5A-5H) releases this repression and promotes cutin biosynthesis, particularly through the activation of *BDG1* (Figure 6E-6G). COP1 has been established as the central regulator of dynamic responses to light-dark cycles, controlling processes such as photomorphogenesis, stomatal development, and conductance (Yang et al., 2005; Liu et al., 2008; Kim et al., 2017b; Lee et al., 2017). The identification of CFLAP1 as a direct target of COP1 provides the missing link explaining how COP1 coordinates diel adaptation in the epidermal cells at both transcriptional and post-translational levels. The dual regulation of cutin biosynthesis and stomatal movement by the COP1-CFLAP1 module suggests a regulatory hub that facilitates protection against water loss, ensuring plant homeostasis across day-night cycles under terrestrial conditions.

Under low-light conditions or darkness, plants exhibit enhanced growth such as hypocotyl elongation and leaf expansion (Apelt et al., 2017). This aerial surface expansion increases the demand for cuticle deposition on newly developed tissues to maintain cellular homeostasis (Ma et al., 2024). The physical properties of cutin support its role as the primary protective layer covering emerging cell surfaces. Lopez-Casado et al. (2007) reported that the cutin matrix displays a low elastic modulus and high strain values relative to the whole cuticle, indicating its flexible and extensible nature. However, the cutin layer alone cannot achieve mechanical stability without the presence of waxes. The incorporation of wax into the cutin matrix converts reversible elastic deformation into irreversible plastic strain, thereby enhancing rigidity. Indeed, the removal of wax from tomato fruit surfaces increased plasticity, indicating that waxes act as structural fillers that reinforce the cutin framework (Petracek and Bukovac, 1995; Domínguez et al., 2011). Such biomechanical characteristics establish a robust cuticle by strengthening the cutin matrix and limiting further surface extension (Khanal et al., 2013). Together, these observations suggest a sequential model in which cutin is initially deposited to protect expanding tissues without impeding cell elongation, followed by wax accumulation that strengthens the cuticle and stabilizes tissue morphology. Considering the provocation of cell elongation during nighttime, concurrent cutin biosynthesis and the following diurnal wax accumulation is critically required. We propose that this mechanism is precisely regulated via COP1-CFLAP1 and LRB-phyB-PIF4 pathways, respectively.

Chronotherapy is the use of circadian rhythm in medicine, which considers environmental and biological rhythms to improve drug treatment efficacy (Selfridge et al., 2016; Kelly et al., 2018). This concept can also be applied to agriculture by aligning genetics and field practices with plants’ daily rhythms (Hotta, 2021). For example, aquaporin expression in roots shows higher abundance during the day, accompanied by stomatal opening (Caldeira et al., 2014; Maurel et al., 2016) which together suggest enhanced daytime water uptake. Our study on diel regulation of cuticle formation may support chronotherapy-inspired crop management. If wax and cutin deposition peaks or off-peaks at predictable time phases, applying pesticides, foliar nutrients, or other external resources at those optimal times may increase uptake efficiency while reducing cost and contamination. Furthermore, breeding or engineering clock and photoreceptor modules to shift the timing of cuticle reinforcement towards periods of maximal drought, heat or UV stress could offer a novel strategy to improve crop resilience and yield.

## MATERIALS AND METHODS

### Plant materials and growth conditions

Arabidopsis mutants and transgenic lines used in this study were listed in Supplemental Table 1. Arabidopsis seeds were sterilized and sown on half-strength Murashige and Skoog (MS) agar plates (pH 5.7) supplemented with 1% sucrose. After stratification for 2 days at 4 °C, plates were placed in growth room under long-day conditions (16 h/8 h, light/dark) at 23 °C and ∼50% humidity. Seven-day-old seedlings were transferred to soil for further experiments. Tobacco (*Nicotiana benthamiana*) were also grown under the same growth conditions as described above.

### Genomic DNA analysis

Genomic DNA was extracted from Arabidopsis seedlings or leaves using extraction buffer (200 mM Tris-HCl pH 7.5, 250 mM NaCl, 25 mM EDTA, 0.5% SDS) and performed PCR to verify the *CFLAP1* locus/transgene using gene-specific primers listed in Supplemental Table S2.

### Gene expression analysis

Total RNA was isolated from 12-day-old seedlings or 3-week-old Arabidopsis leaves using the Total RNA Purification Kit following manufacturer’s instructions (Xenohelix). About 2 μg of total RNA was used for reverse transcription using GoScript™ Reverse Transcriptase (Promega). RT-PCR was performed using 2x Prime Taq Premix (GeNetBio). Real-Time Quantitative Polymerase Chain Reaction (RT-qPCR) was performed using TOPreal™ SYBR Green qPCR PreMIX (Enzynomics) by CFX Opus 96 Real-Time PCR system (Bio-Rad). Transcript levels were normalized to that of *PP2AA3* (At1g13320). The used primers are listed in Supplemental Table 2.

### Dual luciferase reporter assay

Effector constructs were made by insertion of the *CFLAP1* coding sequence (CDS) into pBA002 to express *CaMV35S:MYC-CFLAP1*. The *PIF4* CDS was inserted into pSPYCE vector to generate a *PIF4-eYFPC*. For reporter constructs, the promoter regions of *CYP86A2*, *CYP86A4*, *CYP86A8*, *HTH*, *BDG1*, *GPAT4*, *KAT1*, *KCS2*, *CER1*, and *CER4* were cloned into pGreenII 0800-LUC vector upstream of *LUC* gene. The effector and reporter constructs were introduced into *Agrobacterium tumefaciens* strain GV3101 (pSOUP). *Agrobacterium* strains harboring the effector or reporter constructs were cultured to OD_600_ = 0.8, pelleted, and resuspended in infiltration buffer (10 mM MgCl_2_, 10 mM MES, pH 5.7, 200 μM acetosyringone), adjusted to an OD_600_ of 0.5, and infiltrated into 4-week-old tobacco leaves. Leaf disks were harvested 2 days after infiltration and then dual-luciferase assays were performed using the Dual Luciferase Assay System following the manufacturer’s instructions (Promega).

### The electrophoretic mobility shift assay (EMSA)

To produce MBP-CFLAP1 the *CFLAP1* CDS was cloned into pMAL-c2 vector (NEB). After transformation into *Escherichia coli* (BL21), the cells were grown on in LB media containing 100 μg mL^-1^ ampicillin. Protein expression was induced with 0.1 mM IPTG (isopropyl β-D-1-thiogalactopyranoside) for 3 h at 37 °C. Total proteins were extracted in column buffer (20 mM Tris-HCl pH 7.4, 200 mM NaCl, 1 mM EDTA, 1 mM DTT) from the harvested cells by sonication, purified using amylose resin (NEB), and eluted with column buffer containing 10 mM maltose. Protein concentration and purity were verified using bradford assay (Bio-Rad) and SDS-PAGE (8%). Biotin-labelled single-strand DNA oligonucleotides corresponding to putative cis-elements in the promoter regions of *CYP86A2*, *CYP86A4*, and *BDG1* were synthesized (Bionics) and annealed by heating at 95 °C for 5 min and slowly cooling to RT. EMSA was performed using the LightShift® Chemiluminescent EMSA Kit (Thermo Scientific) following the manufacturer’s protocol with nylon membrane (Hybond). Signal visualization was performed in AI600 Chemidoc Imaging System (GE Healthcare).

### Chromatin immunoprecipitation (ChIP) assay

ChIP assays were performed as described in Gendrel et al., (2005). *35S:MYC-CFLAP1*/Col-0 (14-day-old) and *35S:PIF4-MYC*/Col-0 (12-day-old) seedlings were analyzed in separate experiments, each with *35S:MYC*/Col-0 as a control. Seedlings were crosslinked with 1% formaldehyde under vacuum infiltration for 15 min, and the reaction was quenched with 0.125 M glycine. After washing, samples were frozen, and ground. Nuclei were isolated using sequential extraction buffers, and chromatin was sheared by sonication (Bioruptor® Pico, Diagenode) to an average DNA fragment size of approximately 0.2-1 kb. Immunoprecipitation was conducted using an anti-MYC antibody (Millipore) to capture MYC-, MYC-CFLAP1- or PIF4-MYC-bound chromatin complexes. Following reverse crosslinking and DNA purification, the enrichment of promoter regions of target genes was analyzed by qPCR. The results were normalized to the amplification of the *PP2AA3* gene as an internal control. The primers used in ChIP-qPCR were listed in Supplemental Table 2.

### Luciferase complementation imaging (LCI) assay

The CDSs of *CFLAP1* and *COP1* were cloned into cLUC and nLUC vectors, respectively. *Agrobacterium* harboring these constructs were grown until their OD_600_ reached 1.0, infiltrated into tobacco leaves together with p19 helper strain. After 2 days, the infected leaf areas were infiltrated with luciferin (100 mM) dissolved in 0.1% Triton X-100. After 5 min of dark incubation, the signals were detected in AI600 Chemidoc Imaging System (GE Healthcare).

### Bimolecular fluorescence complementation (BiFC) assay

The CDSs of *CFLAP1, HY5,* and *DEWAX* were cloned into the pSPYCE vector and the *COP1* CDS was cloned into pSPYNE vector. *Agrobacterium* cells were co-infiltrated into tobacco leaves together with the p19 helper strain. After 2 days, the fluorescence signals from tobacco epidermal cells were detected using a confocal laser scanning microscope (TCS SPE, Leica) with a yellow fluorescent protein (YFP) filter (519 nm excitation, 555 nm emission).

### Immunoblot assays

*Agrobacterium* cells carrying *COP1-HA* or *MYC-CFLAP1* were infiltrated into tobacco leaves, treated with 100 µM MG132 8 h, and harvested 24 h after infiltration. Protein was extracted from liquid nitrogen-ground tissues in IP buffer (50 mM Tris-HCl, pH 7.5, 150 mM NaCl, 0.5% NP-40, 1 mM EDTA, 10 mM DTT, 2 mM NaVO_3_, 2 mM NaF, 2 mM PMSF, 3 μg/ml pepstatin A, 3 μg/ml aprotinin, 5 μg/ml leupeptin, 10 μM MG132), separated by 8% SDS-PAGE, transferred to polyvinylidene difluoride (PVDF) membrane (Millipore), and detected using anti-MYC (1:2000, Millipore) or anti-HA (1:4000, Invitrogen) primary antibodies (12 h at 4 °C), and anti-mouse IgG (1:5000, Millipore) secondary antibody (1 h at RT). Signal visualization was performed with Pierce^TM^ ECL Plus Western Blotting Substrate (Thermo Scientific) in AI600 Chemidoc Imaging System (GE Healthcare).

For co-immunoprecipitation (Co-IP) and ubiquitination assays, protein from tobacco leaves co-infiltrated with *Agrobacterium* cells COP1-HA or MYC-CFLAP1 with 100 μM MG132 treatment were extracted in IP buffer. The extracts were incubated with anti-c-MYC conjugated agarose beads (Sigma) on a rotator at 4 °C for 2 h, washed with IP buffer, resuspended in sample buffer (100 mM Tris-HCl, pH 6.8, 4% SDS, 200 mM DTT, 20% glycerol, 0.025% bromophenol blue), and analyzed using anti-MYC (1:2000, Millipore), anti-HA (1:4000, Invitrogen) and anti-Ub (1:2000, Santa Cruz Biotechnology) antibodies.

To assess MYC-CFLAP1 stability, 4-day-old *35S:MYC-CFLAP1*/Col-0 or *35S:MYC-CFLAP1*/*cop1-4* seedlings were incubated in 1/2 MS liquid media with 100 mM cycloheximide (CHX), 100 mM MG132 or DMSO. To assess MYC-CFLAP1 stability over the diurnal cycle, 10-day-old transgenic Arabidopsis seedlings *35S:MYC-CFLAP1*/Col-0 or *35S:MYC-CFLAP1*/*cop1-4* were harvested at indicated time points. Sample was analyzed by western blot with anti-MYC (1:2000, Millipore).

To assess PIF1, PIF3, PIF4, and PIF5 stability, 3-week-old Arabidopsis leaves were harvested at indicated time points. Proteins were extracted using an extraction buffer A (125 mM Tris-HCl, pH 6.8, 4% SDS, 200 mM DTT, 10% glycerol, and protease inhibitors including 2 mM PMSF, 3 μg/ml pepstatin A, 3 μg/ml aprotinin, 5 μg/ml leupeptin, 10 μM MG132) and detected using anti-PIF1, anti-PIF3, anti-PIF4 (1:1000) and anti-PIF5 (1:1000, Agrisera) antibodies. To assess phyB stability, 2-week-old Arabidopsis seedlings were harvested at indicated time points. Proteins were extracted using an extraction buffer B [70 mM Tris-HCl, pH 8.3, 35% ethylene glycol, 98 mM (NH_4_)_2_SO_4_, 7 mM EDTA, 14 mM sodium metabisulfite, 0.07% polyethyleneimine, protease inhibitors (Roche)] and detected using anti-phyB (1:1000, Agrisera) and anti-TCTP antibodies (1:10000).

### TBO Staining

Three-week old Arabidopsis plants were immersed into TBO solution (0.1% TBO, 0.01% Tween-20, Sigma) for 3 minutes, then washed three times in distilled water and photographed using Samsung NX300 Digital camera.

### Cuticular transpiration assay

Three-week-old plants were incubated in darkness for 12 h. The aerial parts were excised and soaked in water for 1 h to equilibrate hydration. After surface water removal, the shoots were weighed every 15 min for up to 3 h using a microbalance in darkness.

### Chlorophyll leaching assay

The aerial parts of three-week-old plants were soaked in 80% ethanol. The extracted chlorophyll amounts according to individual time points (up to 24 h) were displayed as percentages. The amount of extracted chlorophyll was quantified by measuring the absorbance at 647 and 664 nm using a diode array spectrophotometer (Ultrospec 3100 pro; Amersham Biosciences).

### TEM analysis

3-week-old rosette leaves from Col-0 and *cop1-4* were harvested which were subsequently fixed and analyzed as previously described (Kim et al., 2025). Shortly, the leaves were fixed o/n at 4°C using Karnovsky’s fixation solution (2% paraformaldehyde, 2.5% glutaraldehyde, 0.1 M sodium cacodylate buffer (pH 7.4)) and post-fixed using 1% osmium tetroxide (OsO_4_) at 4°C for 1 hour. Samples were dehydrated through a graded ethanol series and gradually embedded with Spurr’s epoxy resin (medium hardness, Ted Pella) with sequential exchange of EtOH:resin=2:1, 1:1, 1:2, and finally 100% resin o/n. Samples were hardened for 36∼48 hours at 60 ℃. Resin-embedded samples were sectioned into 80–100 nm slices using an ultramicrotome (RMC Products), placed on a grid and stained with uranyl acetate and lead citrate prior to observation with TEM (Jeol, JEM-2100F).

### Cuticular wax and cutin analysis

Three-week-old Arabidopsis leaves were used for cuticular wax and cutin analyses. Cuticular waxes were extracted by 4 ml chloroform for 30 sec. Internal standards (2 μg *n*-Octacosane, 1 μg 1-tricosanol, and 2 μg heptadecanoic acid; Sigma) were added to the extracts, evaporated under a stream of nitrogen gas, and trimethylsilylated by heating with 100 μl of N, O-Bis (trimethylsilyl) trifluoroacetamide (Sigma) and 100 μl of pyridine (Sigma) at 100 °C for 30 min. After evaporation under nitrogen, samples were dissolved in heptane:toluene (1:1, v/v). Wax compounds were quantified using GC-FID (GC-2010 Plus, Shimazu) using DB-5 (60 m) column (Agilent) as previously described (Kim et al., 2021). For cutin analysis, leaves were delipidated and depolymerized. C17:0 FAME and ω-pentadecalactone (Sigma) were used as internal standards, as previously described (Lee et al., 2019). Derivatization of cutin monomers was performed with 100 μl of pyridine (Sigma) and 100 μl of acetic anhydride (J.T.Baker) at 60 °C for 2 h. Solvent was evaporated under nitrogen gas and redissolved in heptane/toluene (1:1, v/v). Cutin monomers were quantified using the same equipment described above. The oven temperature increased to 300°C at a rate of 2.5 °C/min and maintained at 300 °C for 3 min.

## AUTHOR CONTRIBUTIONS

Q.H.D., H.J.K., J.C., and M.C.S. conceived the study. Q.H.D., H.J.K., D.-M.C., and S.-H.K. performed the experiments and jointly analyzed the data with J.C., J.-I.K., and M.C.S. Q.H.D., H.J.K., J.C., and M.C.S. wrote the manuscript, and J.-I.K. contributed to scientific discussions and provided a critical review of the manuscript.

## ACKNOWLEDGEMENTS

The authors thank Takato Imaizumi (University of Washington) and Woe-Yeon Kim (Gyeongsang National University) for sharing Arabidopsis mutant seeds listed in Supplemental Table 1. The authors are grateful to Ryeo Jin Kim for TEM analysis.

## SUPPLEMENTAL INFORMATION

The following materials are available in the online version of this article.

**Supplemental Figure S1.** Cuticular wax composition and amounts from wild type (Col-0 /Ler), phytochrome mutants, and a *phyB* overexpressing line (*PHYB OX*).

**Supplemental Figure S2.** Expression patterns of *PIFs* transcripts and proteins in Arabidopsis WT during the daytime and nighttime

**Supplemental Figure S3.** Cuticular wax composition and amounts from wild type (Col-0), *pif4*, *pifQ*, and *PIF4/pifQ,* and *PIF4 OX*.

**Supplemental Figure S4.** Cutin monomer composition and amounts from wild type (Col-0)*, pif4, pifQ,* and *PIF4/pifQ* leaves.

**Supplemental Figure S5.** Cuticular wax composition and amounts from wild type (Col-0) and *lrb123*.

**Supplemental Figure S6.** Isolation of the *cflap1* mutant.

**Supplemental Figure S7.** Cutin monomer and cuticular wax composition and amounts from wild type (Col-0)*, cflap1-1,* and *cflap1-2* leaves.

**Supplemental Figure S8.** Expression patterns of *CFLAP1* transcripts and proteins in Arabidopsis WT and *CFLAP1 OX*/Col-0 during the daytime and nighttime

**Supplemental Figure S9.** IPTG induction test and purification of recombinant MBP-CFLAP1 protein

**Supplemental Figure S10.** Cutin monomer and cuticular wax composition and amounts from wild type (Col-0)*, cop1-4,* and *cop1-6* leaves.

**Supplemental Figure S11.** Genotyping of *cop1-4, cflap1-1, cflap1-2,* and *cflap1cop1-4*.

**Supplemental Figure S12.** Cutin monomer and cuticular wax composition and amounts from wild type (Col-0)*, cflap1-1, cflap1-2, cop1-4, cflap1-1 cop1-4* and *cflap1-2 cop1-4* leaves.

**Supplemental Table S1.** The list of Arabidopsis mutant and transgenic lines used in this study.

**Supplemental Table S2.** Oligonucleotide primers used in this study.

