## Supplemental Figures for "Unraveling diel regulation of cuticle biosynthesis"

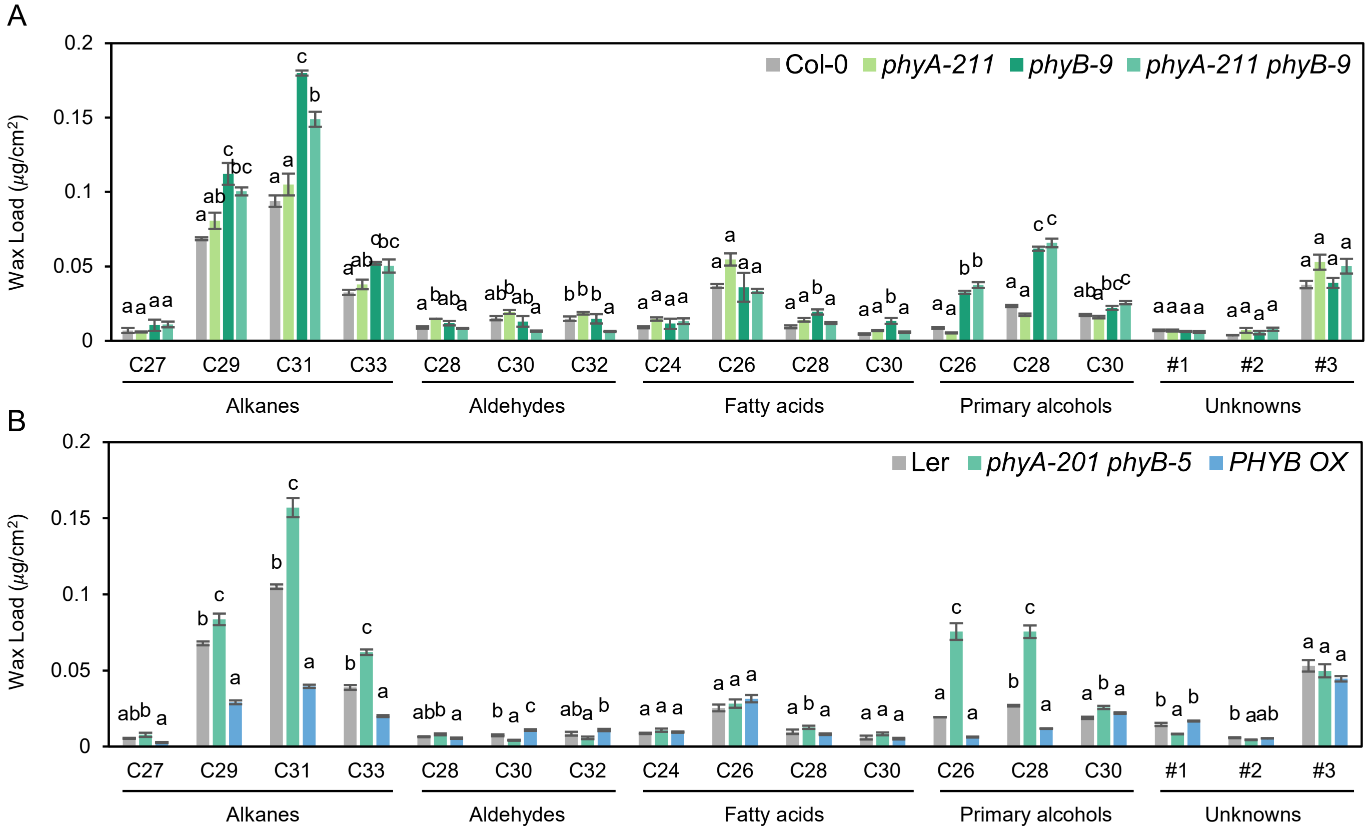


**Figure S1. Cuticular wax composition and amounts from wild type (Col-0 /Ler), phytochrome mutants, and a *phyB* overexpressing line (*PHYB OX*).**

(A) Cuticular waxes from 3-week-old wild-type (Col-0), *phyA-211*, *phyB-9*, and *phyA-211 phyB-9* leaves analyzed using GC-FID. (B) Cuticular waxes from 3-week-old wild-type (Ler), *phyA-201 phyB-5*, and *PHYB OX* leaves analyzed using GC-FID. (A and B) Values indicate mean ± SD from three replicate experiments. Different letters indicate statistically significant differences using one-way ANOVA with Tukey's test ($P < 0.01$).

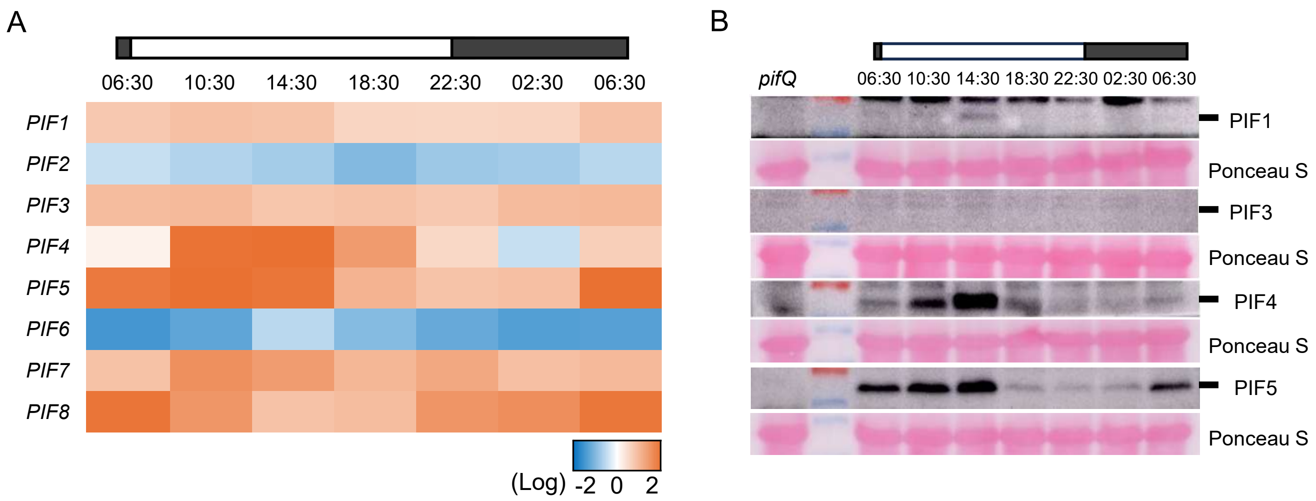


**Figure S2. Expression patterns of *PIFs* transcripts and proteins in Arabidopsis WT during the daytime and nighttime**

(A) Expression profiles of *PIF1-8* in wild-type (Col-0). Transcript levels were examined by RT–qPCR. Transcript abundance is shown on a log scale, CT threshold of 28 used to demarcate high from low transcript levels. Data represent the average of three replicates. (B) Diurnal patterns of PIF1, PIF3, PIF4, and PIF5 protein abundances in Col-0. PIF1, PIF3, PIF4, and PIF5 protein levels were analyzed by immunoblot using anti-PIF1, anti-PIF3, anti-PIF4, and anti-PIF5 antibodies, respectively. *pifQ* harvested at 06:30 was used as a negative control. Total protein loading was assessed by Ponceau S staining. (A and B) 3-week-old leaves of wild type (Col-0) were harvested at indicated time points.

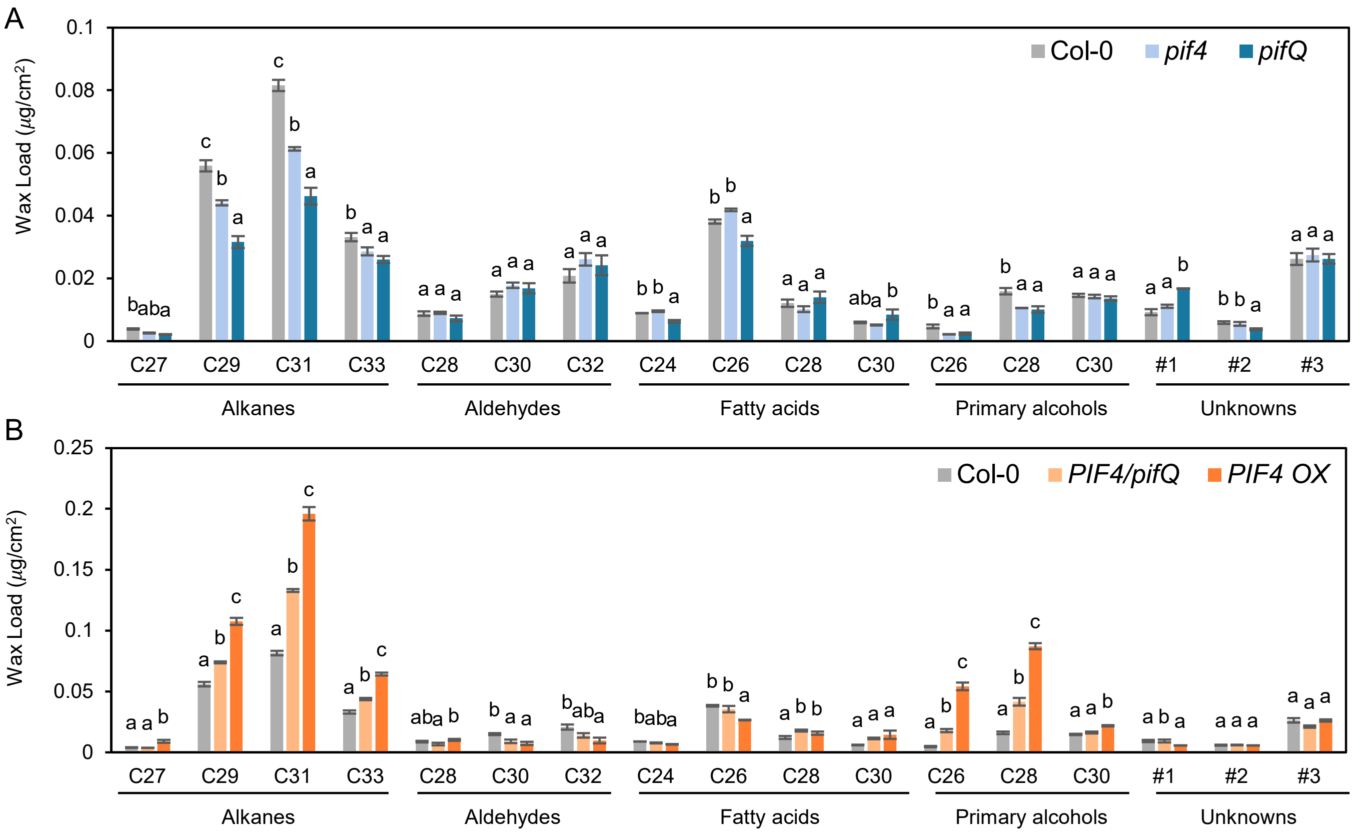


**Figure S3. Cuticular wax composition and amounts from wild type (Col-0), *pif4*, *pifQ*, and *PIF4/pifQ*, and *PIF4 OX*.**

(A) Cuticular waxes from 3-week-old Col-0, *pif4*, and *pifQ* leaves analyzed using GC-FID. (B) Cuticular waxes from 3-week-old Col-0, *PIF4pro:PIF4-MYC/pifQ* (*PIF4/pifQ*), and *35Spro:PIF4-MYC*/Col-0 (*PIF4 OX*) analyzed using GC-FID. (A and B) Error bars indicate ±SD from three replicate experiments. Different letters indicate statistically significant differences using one-way ANOVA with Tukey's test ($P < 0.01$).

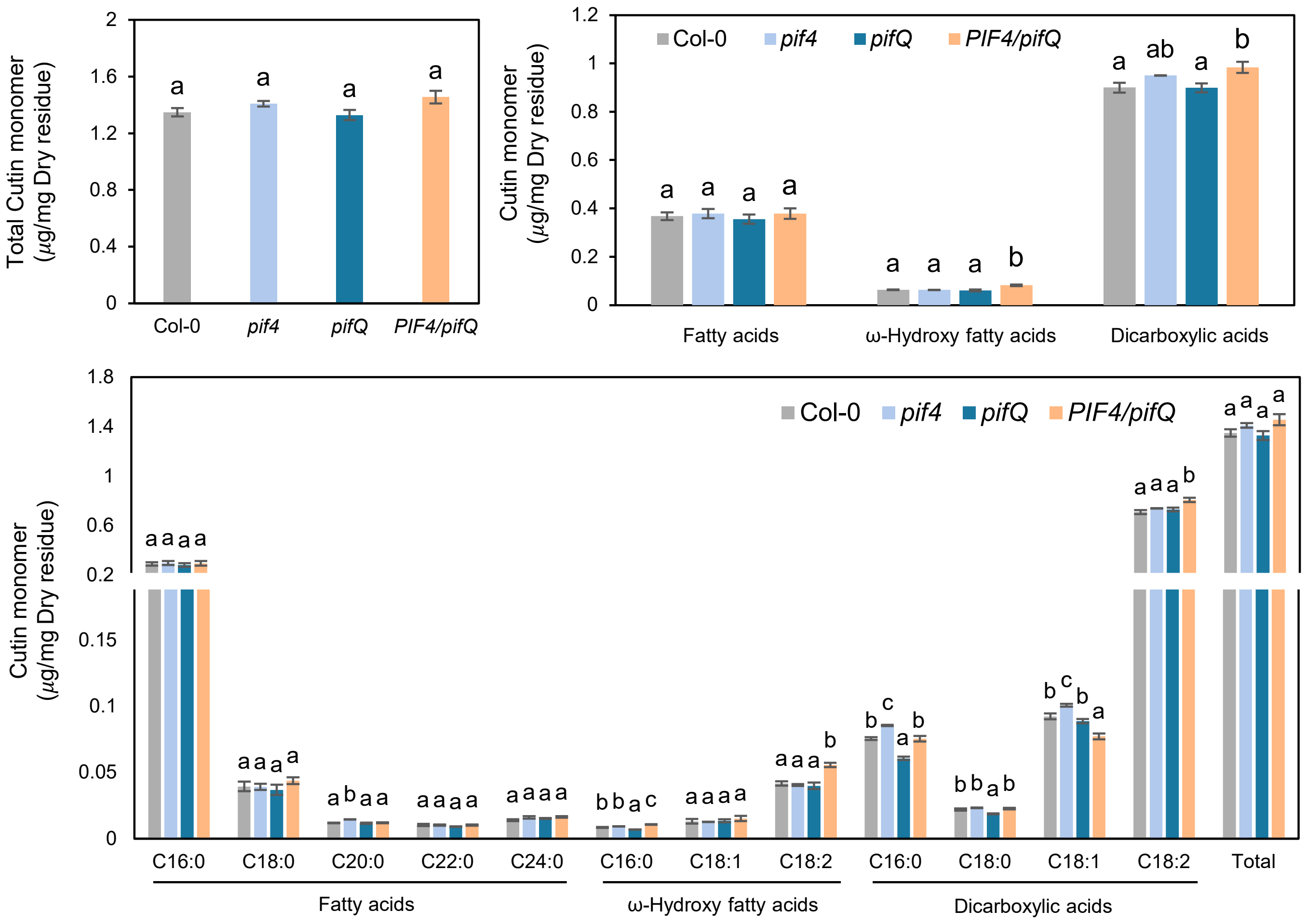


**Figure S4. Cutin monomer composition and amounts from wild type (Col-0), *pif4, pifQ,* and *PIF4/pifQ* leaves.**

Cutin monomers from 3-week-old Col-0, *pif4*, *pifQ,* and *PIF4/pifQ* leaves analyzed using GC-FID. Values represent mean ±SD from three replicate experiments. Different letters indicate statistically significant differences using one-way ANOVA with Tukey's test ($P < 0.01$).

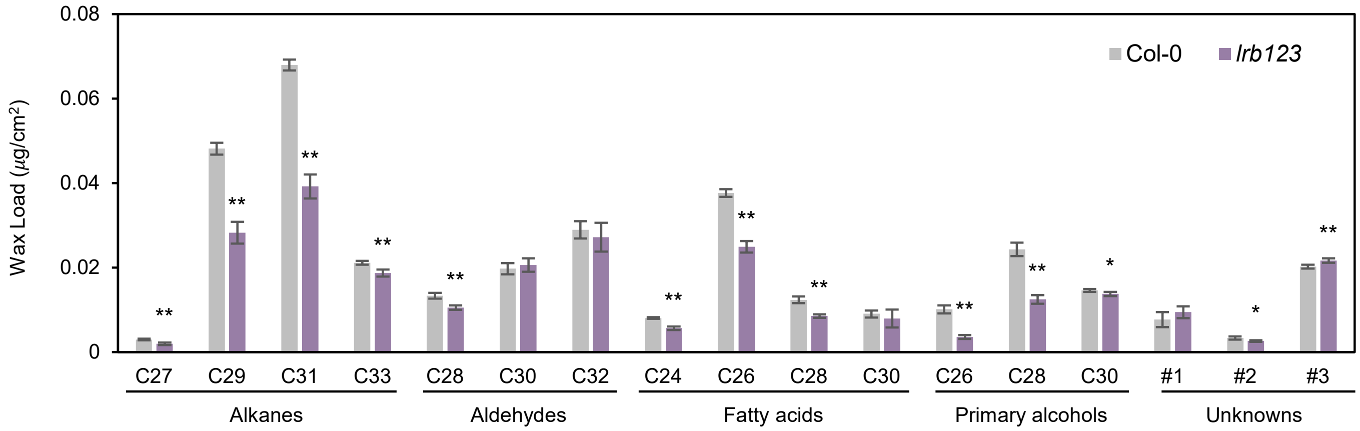


**Figure S5. Cuticular wax composition and amounts from wild type (Col-0) and *lrb123*.** Cuticular waxes from 3-week-old Col-0 and *lrb123* leaves analyzed using GC-FID. Values represent mean ± SD from three replicate experiments. Asterisks indicate statistically significant differences determined by Student's t-test (*, $P < 0.05$; **, $P < 0.01$).

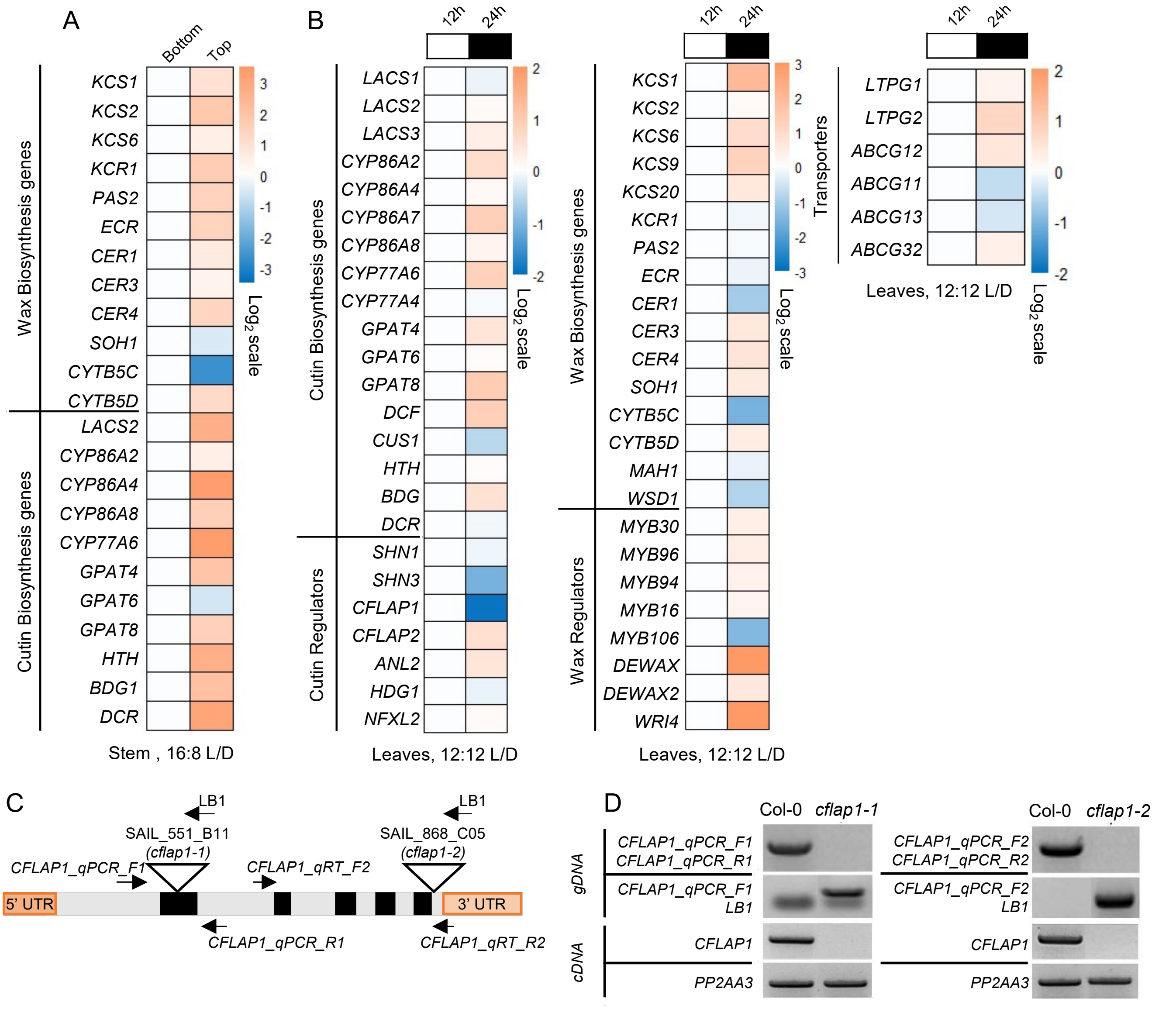


**Figure S6. Isolation of the *cflap1* mutant.**
(A and B) Microarray heatmaps of genes involved in cuticular wax and cutin biosynthesis, transport and regulation. (C) Schematic representation of *CFLAP1* showing T-DNA insertions at 2 chromosomal sites and position of used primers. (D) Genomic DNA PCR and RT-PCR analysis of Col-0, *cflap1-1,* and *cflap1-2* seedlings. *PP2AA3* (At1g13320) was used as a reference gene. LB, Left border; UTR, untranslated region; L/D, Light/Dark.

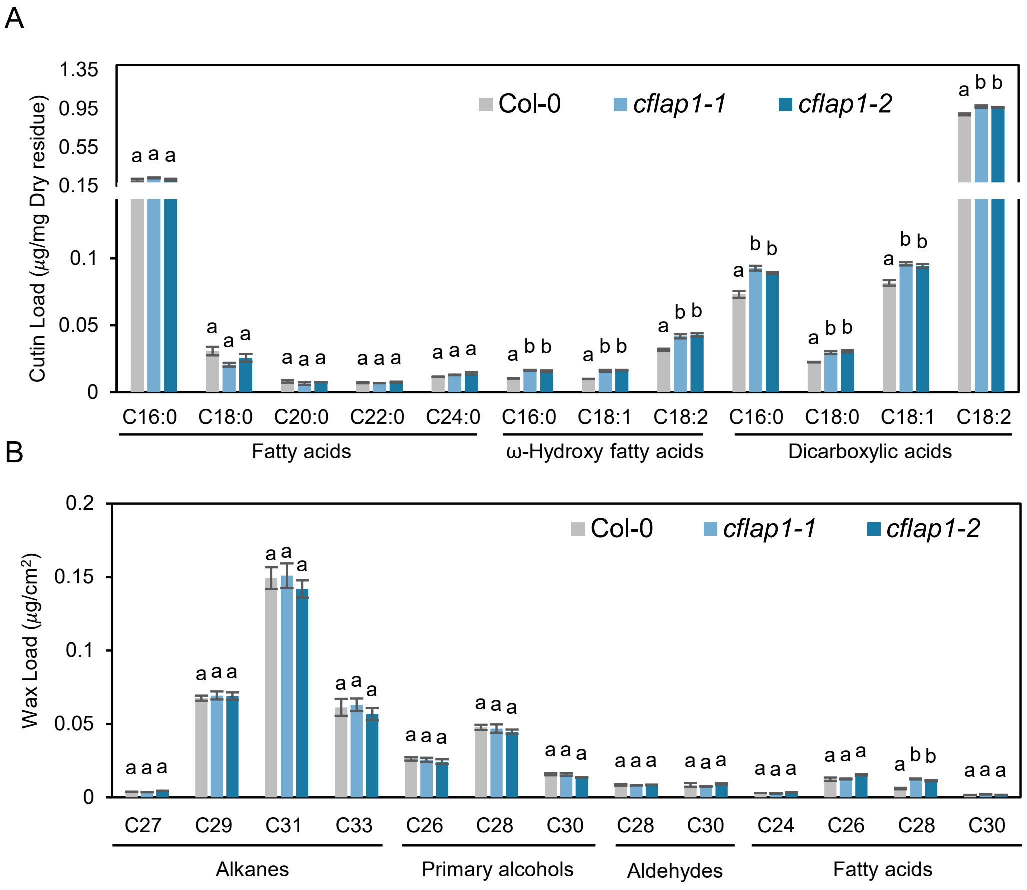


**Figure S7. Cutin monomer and cuticular wax composition and amounts from wild type (Col-0), *cflap1-1*, and *cflap1-2* leaves.**

(A and B) Cutin monomers (A) and cuticular waxes (B) from 3-week-old Col-0, *cflap1-1*, and *cflap1-2* leaves analyzed using GC-FID. Each value represents the mean ± SD of three individual replicates. Different letters indicate statistically significant differences using one-way ANOVA with Tukey's test ($P < 0.01$). c

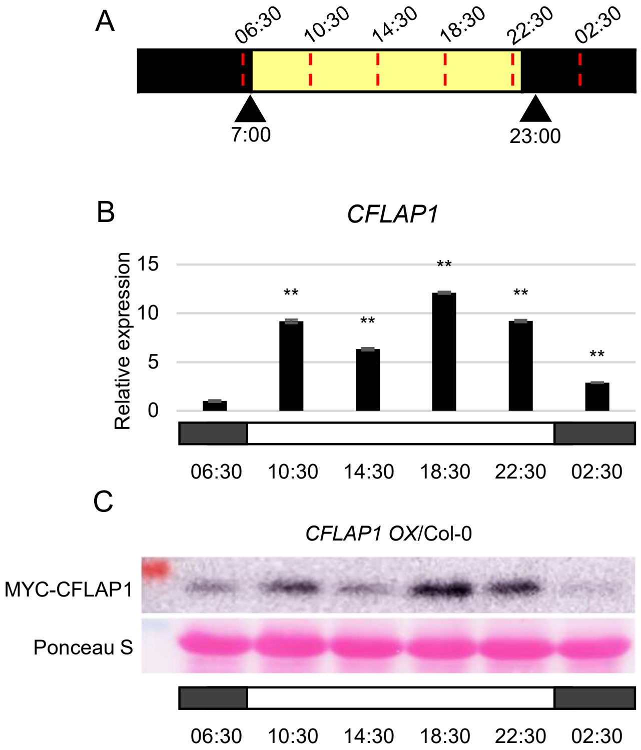


**Figure S8. Expression patterns of *CFLAP1* transcripts and proteins in Arabidopsis WT and *CFLAP1 OX*/Col-0 during the daytime and nighttime**

(A) Schematic of the 16-h light/8-h dark photoperiod used for 12-day-old seedlings. Yellow and black bar indicates the light and dark phases, respectively. Red dashed lines mark the time points at which samples were collected. (B) Diurnal change of *CFLAP1* transcript levels in wild-type (Col-0). Transcript levels were normalized to *PP2AA3* (At1g13320). Asterisks indicate statistically significant differences determined by Student's t-test (*, $P < 0.05$; **, $P < 0.01$). (C) Diurnal change of CFLAP1 protein abundances in *CFLAP1 OX*/Col-0 seedlings. Ponceau S staining shows protein loading controls.

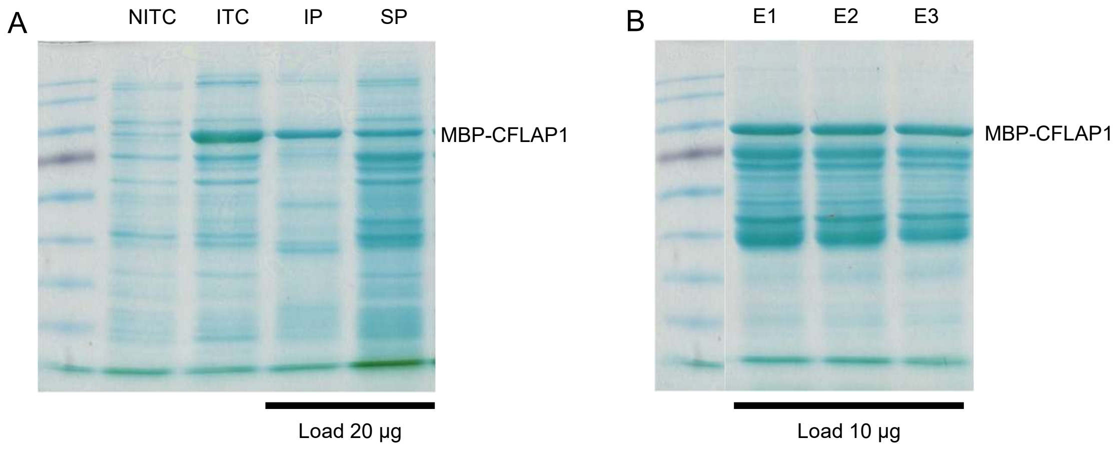


**Figure S9. IPTG induction test and purification of recombinant MBP-CFLAP1 protein**
(A) IPTG (isopropyl β-D-1-thiogalactopyranoside) induction test of *E. coli* cells harboring *MBP-CFLAP1* construct. Lane 1, non-induced total cell (NITC); Lane 2, induced total cell (ITC) with 0.1 mM IPTG at 37 $^{o}$C for 3 h; Lane 3, insoluble protein (IP); Lane 4, soluble protein (SP). (B) Purification of MBP-CFLAP1 from *E. coli* cells. Lane 1 to Lane 3 (E, elution) are protein eluents from amylose resin.

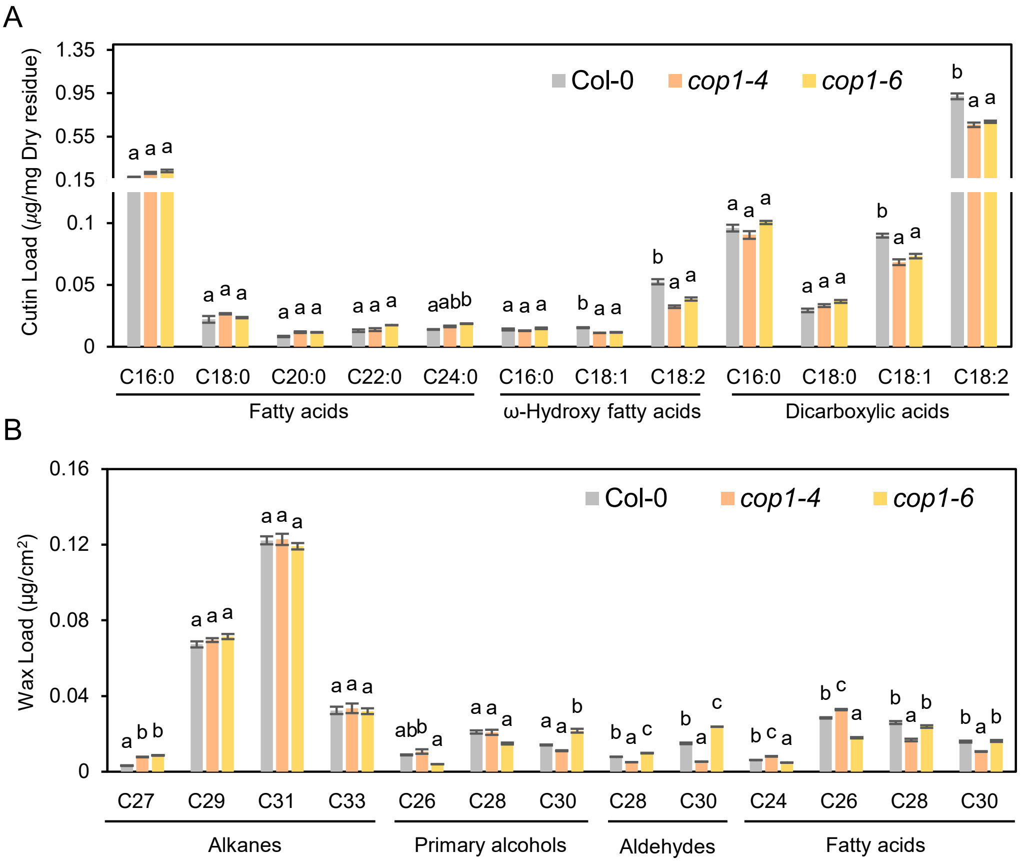


**Figure S10. Cutin monomer and cuticular wax composition and amounts from wild type (Col-0), *cop1-4,* and *cop1-6* leaves.**

(A and B) Cutin monomers (A) and cuticular waxes (B) from 3-week-old Col-0, *cop1-4*, and *cop1-6* leaves analyzed using GC-FID. Each value represents the mean ±SD of three individual replicates. Different letters indicate statistically significant differences using one-way ANOVA with Tukey's test ($P < 0.01$).

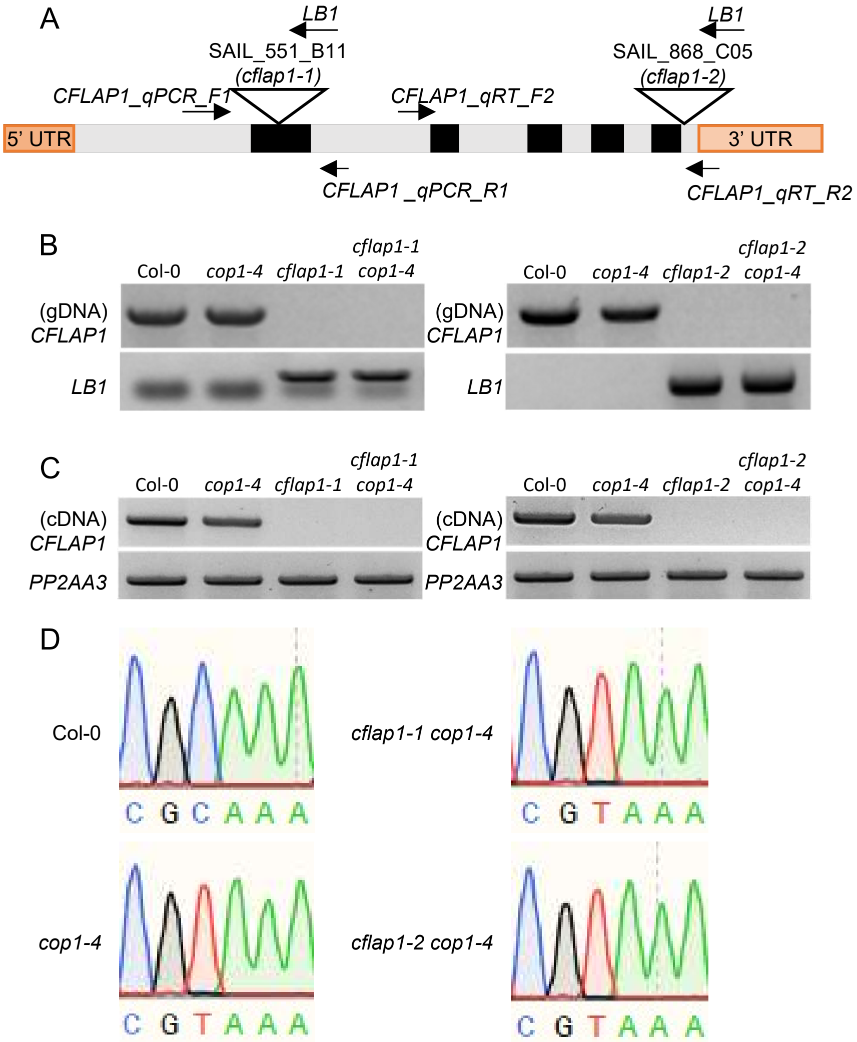


**Figure S11. Genotyping of *cop1-4, cflap1-1, cflap1-2,* and *cflap1 cop1-4.***
(A) Schematic representation of *CFLAP1* showing T-DNA insertions at 2 chromosomal sites and used primers. (B and C) Genotyping of wild-type (Col-0), *cflap1-1, cflap1-2, cop1-4, cflap1-1 cop1-4,* and *cflap1-2 cop1-4* by genomic DNA PCR and RT-PCR analysis. *PP2AA3 (*At1g13320) was used as a reference gene. (D) Sequencing chromatograms of Col-0, *cop1-4, cflap1-1 cop1-4,* and *cflap1-2 cop1-4* seedlings. LB, Left border; UTR, Untranslated region.

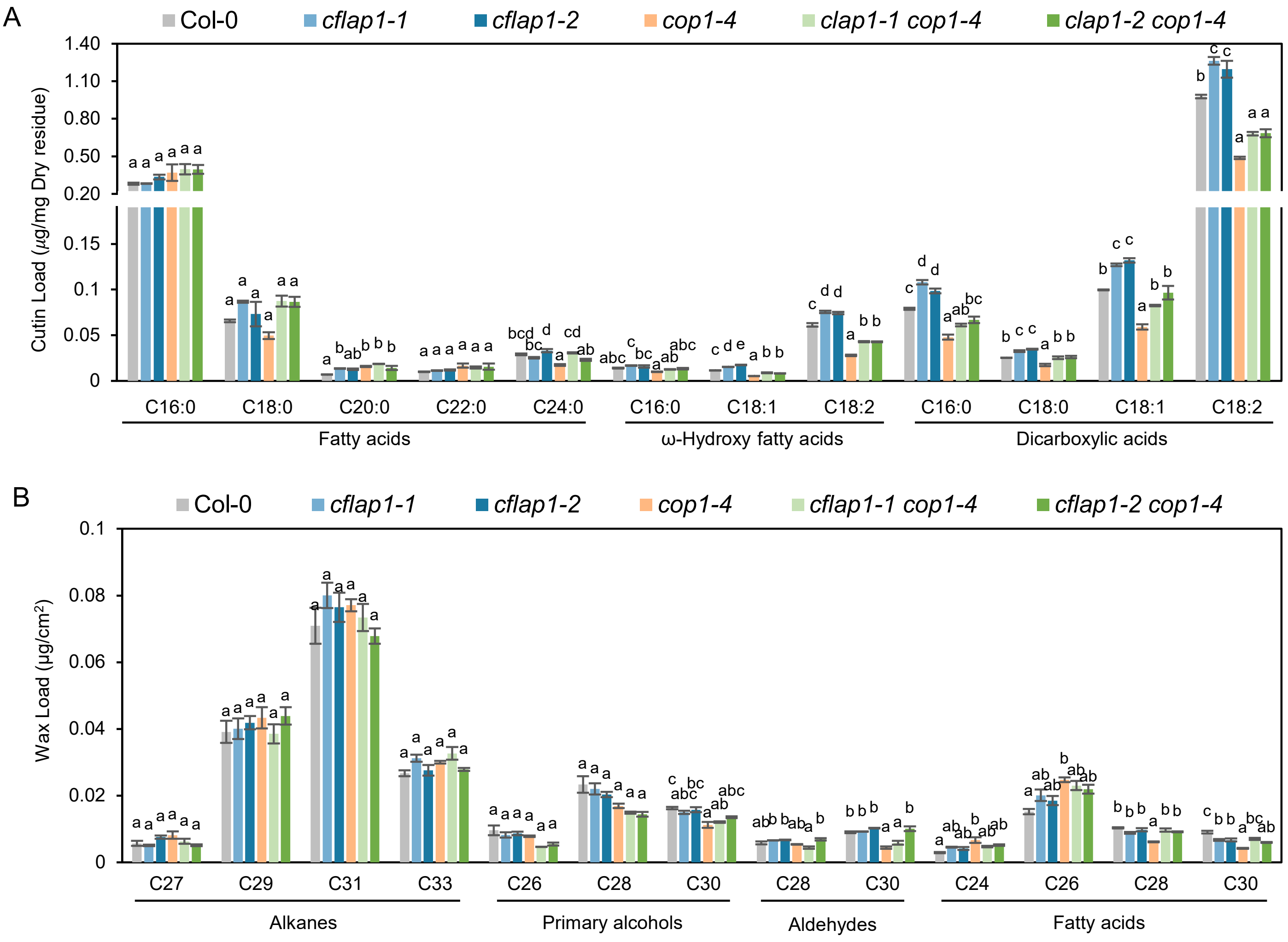


**Figure S12. Cutin monomer and cuticular wax composition and amounts from wild type (Col-0), *cflap1-1, cflap1-2, cop1-4, cflap1-1 cop1-4* and *cflap1-2 cop1-4* leaves.**

(A and B) Cutin monomers (A) and cuticular waxes (B) from 3-week-old Col-0, *cflap1-1, cflap1-2, cflap1-1 cop1-4,* and *cflap1-2 cop1-4* leaves analyzed using GC-FID. Each value represents the mean ±SD of three individual replicates. Different letters indicate statistically significant differences using one-way ANOVA with Tukey's test ($P < 0.01$).
